# Genome streamlining, proteorhodopsin, and organic nitrogen metabolism in freshwater nitrifiers

**DOI:** 10.1101/2021.01.19.427344

**Authors:** Justin C. Podowski, Sara F. Paver, Ryan J. Newton, Maureen L. Coleman

## Abstract

Microbial nitrification is a critical process governing nitrogen availability in aquatic systems. Freshwater nitrifiers have received little attention, leaving many unanswered questions about their taxonomic distribution, functional potential, and ecological interactions. Here, we reconstructed genomes to infer the metabolism and ecology of free-living picoplanktonic nitrifiers across the Laurentian Great Lakes, a connected series of five of Earth’s largest lakes. Surprisingly, ammonia oxidizing Bacteria (AOB) related to *Nitrosospira* dominated over ammonia oxidizing Archaea (AOA) at nearly all stations, with distinct ecotypes prevailing in the transparent, oligotrophic upper lakes compared to Lakes Erie and Ontario. Unexpectedly, one ecotype of *Nitrosospira* encodes proteorhodopsin, which could enhance survival in conditions where ammonia oxidation is inhibited or substrate limited. Nitrite oxidizing Bacteria (NOB) *Ca.* Nitrotoga and *Nitrospira* fluctuated in dominance, with the latter prevailing in deeper, less productive basins. Genome reconstructions reveal highly reduced genomes and features consistent with genome streamlining, along with diverse adaptations to sunlight and oxidative stress and widespread capacity for organic nitrogen use. Our findings expand the known functional diversity of nitrifiers and establish their ecological genomics in large lake ecosystems. By elucidating links between microbial biodiversity and biogeochemical cycling, our work also informs ecosystem models of the Laurentian Great Lakes, a critical freshwater resource experiencing rapid environmental change.

**Importance:** Microorganisms play critical roles in Earth’s nitrogen cycle. In lakes, microorganisms called nitrifiers derive energy from reduced nitrogen compounds. In doing so, they transform nitrogen into a form that can ultimately be lost to the atmosphere by a process called denitrification, which helps mitigate nitrogen pollution from fertilizer runoff and sewage. Despite their importance, freshwater nitrifiers are virtually unexplored. To understand their diversity and function, we reconstructed genomes of freshwater nitrifiers across some of Earth’s largest freshwater lakes, the Laurentian Great Lakes. We discovered several new species of nitrifiers specialized for clear low nutrient waters, and distinct species in comparatively turbid Lake Erie. Surprisingly, one species may be able to harness light energy using a protein called proteorhodopsin, despite the fact that nitrifiers typically live in deep dark water. Our work reveals unique biodiversity of the Great Lakes and fills key gaps in our knowledge of an important microbial group, the nitrifiers.

## Introduction

The oxidation of ammonia to nitrate powers the growth of nitrifying microorganisms and represents a critical flux in the global nitrogen cycle. Microbial nitrification of ammonia released from organic matter degradation produces nitrate, which can then be removed from the system by denitrification (1). As chemolithoautotrophs, nitrifiers are also a major source of dark carbon fixation (2), which may contribute significant organic carbon to the microbial food web of the ocean’s interior (3–5) and of deep freshwater lakes (6).

Microbial nitrifiers are found in Archaea and several phyla of Bacteria, spanning diverse physiology and ecology; understanding the drivers and consequences of this diversity across ecosystems is a fundamental challenge. Ammonia-oxidizing Archaea (AOA) in the phylum *Thaumarchaeota* dominate the mesopelagic oceans (7), likely due to their high affinity for ammonia (8) and streamlined genomes (9). In freshwater systems, AOA are abundant in some oligotrophic lakes, while ammonia-oxidizing Bacteria (AOB) affiliated with the *Nitrosomonadaceae* (Betaproteobacteria) are highly variable but tend to dominate more eutrophic systems (10–16). Complicating this picture, however, there is considerable physiological variation within each group, such as low-nutrient-adapted clades of AOB (17, 18) and the ability of some strains to use alternative substrates like urea (18, 19). Within the AOA, there are also distinct ecotypes that appear to segregate with depth in the water column, in both marine (7) and freshwater systems (10). In freshwaters especially — which are poorly characterized compared to the oceans — it remains difficult to predict which AOA and AOB taxa are likely to dominate in a given system (16).

For aquatic nitrite oxidizers, which span the phyla *Nitrospira, Nitrospinae,* and *Proteobacteria,* niche differentiation is even less clear. The oceans are dominated by exclusively marine lineages (2, 20), consistent with ancient salinity-associated divergence. Cultivated strains of NOB show variation in substrate affinity and physiology (20–22), but the phylogenetic conservation of these traits, and their influence on environmental distributions, are poorly understood. Moreover, recent studies have discovered that NOB are capable of alternative energy metabolisms (23, 24) and can access nitrogen from cyanate and urea (25, 26), expanding their ecological potential. In freshwater systems, the NOB *Ca.* Nitrotoga (Betaproteobacteria) was only recently discovered to be widespread (27), and the diversity of this genus and factors favoring its success are unknown.

Here, we use the Laurentian Great Lakes as a model system to examine niche partitioning among planktonic freshwater nitrifiers. The Great Lakes hold 20% of Earth’s surface freshwater and more than half of this volume receives little to no light (< 1% surface irradiance). This system, while hydrologically connected, spans strong trophic and chemical gradients: ultraoligotrophic Lake Superior supports rates of primary production and nitrification comparable to the ocean gyres (28, 29), while Lake Erie supports greater production (30) and more than 70- fold higher nitrification rate (31). Between these extremes, Lake Ontario has low ambient ammonium concentrations like Lake Superior (32) but nitrification rates up to four times higher (33). While previous studies reported that AOA and AOB dominate Lakes Superior and Erie, respectively (14, 29), recent community profiling has revealed broader diversity in both ammonia-oxidizing and nitrite-oxidizing lineages (34–36). We sought to link taxonomic, genomic, and metabolic diversity of nitrifiers with the varied biogeochemistry of the Great Lakes, using genome reconstructions and abundance profiling. Our results uncover novel lineages and metabolic capabilities, and provide the first large-scale assessment of freshwater nitrifier genomics.

## Results and Discussion

### Niche partitioning of nitrifiers across the Great Lakes

To map free-living picoplanktonic (here defined as cells that pass through a 1.6µm filter) nitrifiers across the Great Lakes, we searched our recent 16S rRNA datasets for known nitrifying taxa (34). We detected putative AOB in the genus *Nitrosospira* (Betaproteobacteria, family *Nitrosomonadaceae*) and AOA similar to *Nitrosarchaeum* (family *Nitrosopumilaceae*), along with putative NOB in the genera *Ca.* Nitrotoga (Betaproteobacteria, family *Gallionellaceae*) and *Nitrospira* (family *Nitrospiraceae*). We did not detect 16S rRNA amplicons from *Nitrosococcus*, *Nitrococcus*, *Nitrospina* or *Nitrobacter*. The highest relative abundances of picoplanktonic nitrifiers were observed in deep samples from eastern Erie and Ontario (9-24% of total amplicons), compared to 2-14% in Michigan, Huron and Superior; Erie and Ontario also have higher cell concentrations and higher surface chlorophyll (Dataset S1). The relative abundance of nitrifiers was negatively correlated with photosynthetically active radiation (PAR; Spearman’s rho = -0.89, p < 2.2e-16) and reached a maximum below the depth of 1% PAR in each lake, up to 20% of amplicon sequences (Fig S1a). The relative abundances of ammonia- and nitrite-oxidizing taxa were strongly correlated (Spearman’s rho = 0.918, p < 2.2e-16; Fig S1b). Free-living nitrifiers were rare (<0.1% relative abundance) in bottom water samples from the southern basin of Huron (HU15M) and western basin of Erie (HU91M); these two stations are the shallowest in our dataset and have relatively high light penetration to the bottom (∼1% PAR). Chlorophyll-*a* concentration was also negatively correlated with the relative abundance of nitrifiers (Spearman’s rho = -0.677, p < 1.7e-7; Fig S1c). These findings are consistent with previous work demonstrating photoinhibition of nitrification (37–40), as well as potential competition with phototrophs for ammonium (41).

The taxonomic assemblage of nitrifiers differed across lakes and even among stations within a lake (Fig 1, Dataset S1), in association with variable productivity and nitrogen availability. Surface ammonium is typically below 300 nM except in Erie, where it is several-fold higher and spatially variable; nitrate, on the other hand, is very high across the lakes but lowest in Erie due to biological uptake (42, 43). Few measurements of urea exist but it can exceed ammonium (44) (Dataset S2). AOB (*Nitrosomonadaceae*) were observed across all lakes. In contrast, AOA (*Nitrosopumilaceae*) sequences only exceeded 0.5% relative abundance at the three deepest stations (SU08M, MI41M, ON55M) where the ratio of AOB:AOA ranged from 10:1 to 1:3. We found pronounced shifts in the dominant NOB across stations (Fig 1), and all stations except those in Lake Ontario showed strong dominance (greater than ten-fold) of either *Ca.* Nitrotoga (family *Gallionellaceae*) or *Nitrospira*. *Nitrospira* was the only nitrite oxidizer detected in Superior and the dominant nitrite oxidizer in parts of Michigan (MI41M, MI18M). By contrast, *Ca.* Nitrotoga was the only nitrite oxidizer observed in Erie and Huron and the dominant nitrite oxidizer at the shallowest station in Michigan (MI27M). Within each taxon, a single 16S rRNA oligotype dominated the AOA, *Ca*. Nitrotoga, and *Nitrospira,* while several oligotypes of *Nitrosomonadaceae* shifted abundance across samples (Fig S2), consistent with ecotypic diversity as discussed below.

**Figure 1.**
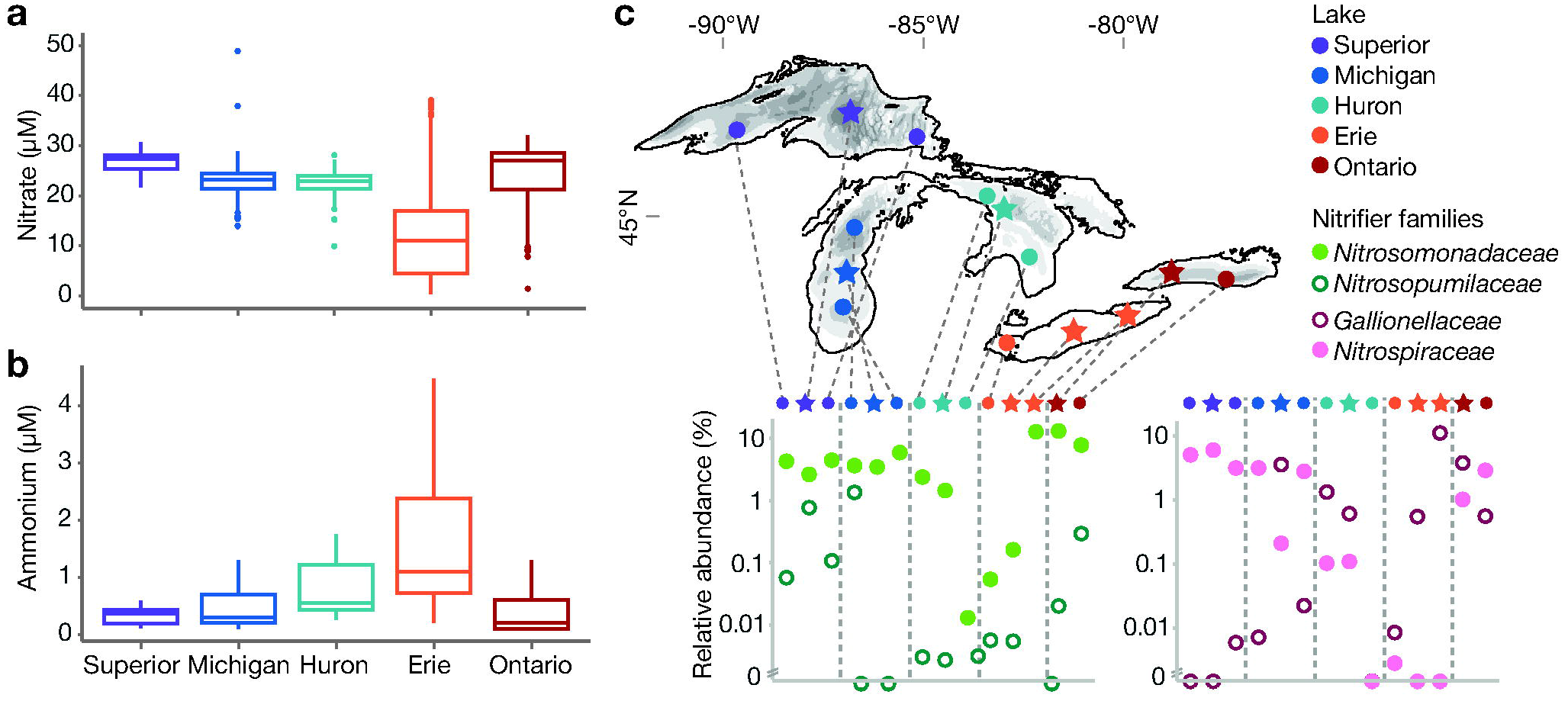
Dissolved inorganic nitrogen availability and distribution of nitrifiers across the Great Lakes. **(a)** Oxidized nitrogen concentrations. Values include NOx concentrations from published studies (n=128; (14, 33, 35, 137, 138)), US EPA Water Quality Surveys in 2012 and 2013 (n=1626 from GLENDA database), and this study (n=20). **(b)** Ammonium concentrations. Values are derived from the literature as in (a) (n=118) and from this study (n=20). **(c)** Distribution of nitrifiers across the Great Lakes. Top panel: map of sampling stations; stars indicate stations chosen for metagenome analysis. Bottom panel: relative abundance of ammonia-oxidizing (green) and nitrite-oxidizing (pink) families based on 16S rRNA V4-V5 amplicon sequencing, sampled in the mid-hypolimnion (except western Erie, sampled 1m from bottom). Data is plotted roughly west to east as indicated on the map.

### Ecotypic variation in abundant streamlined Nitrosospira

We reconstructed 15 genomes of the AOB *Nitrosospira*, substantially expanding genome descriptions for this genus (45–47). Based on a phylogenomic tree, free-living Great Lakes *Nitrosospira* fall into two major clades, both of which are distinct from published species; each of these clades also includes MAGs recovered from lakes Biwa and Baikal, suggesting novel globally distributed freshwater lineages (Fig S3). One clade, which we call NspGL1, has a highly reduced genome (median 1.42 Mb) and low G+C content (40.7%) (Fig 2, Dataset S3). The second clade was resolved into three subclades (denoted NspGL2a, 2b, and 3; Fig S3) based on phylogeny and average nucleotide identity (ANI), all with small genome sizes of 1.45-1.68 Mb and 50% G+C content (Fig 2, Dataset S3). Compared to 86 reference *Nitrosomonadaceae* genomes, Great Lakes *Nitrosospira* genomes are not only smaller (median estimated complete genome size: reference = 3.21 Mb, GL = 1.45 Mb; Table 1), but also have shorter intergenic spacers, fewer paralogs, fewer pseudogenes, and fewer sigma factors (Table 1, Fig S4, Dataset S4), consistent with genome streamlining to reduce resource demands (48). Based on short read mapping, these subclades are ecologically distinct: NspGL1 and NspGL2b — with the smallest genomes — are the dominant AOB in the upper oligotrophic lakes, while NspGL2a is only abundant in Ontario and NspGL3 is only abundant in Erie (Fig 2). Hereafter we refer to these subclades as ecotypes due to their phylogenetic and ecological divergence.

**Figure 2.**
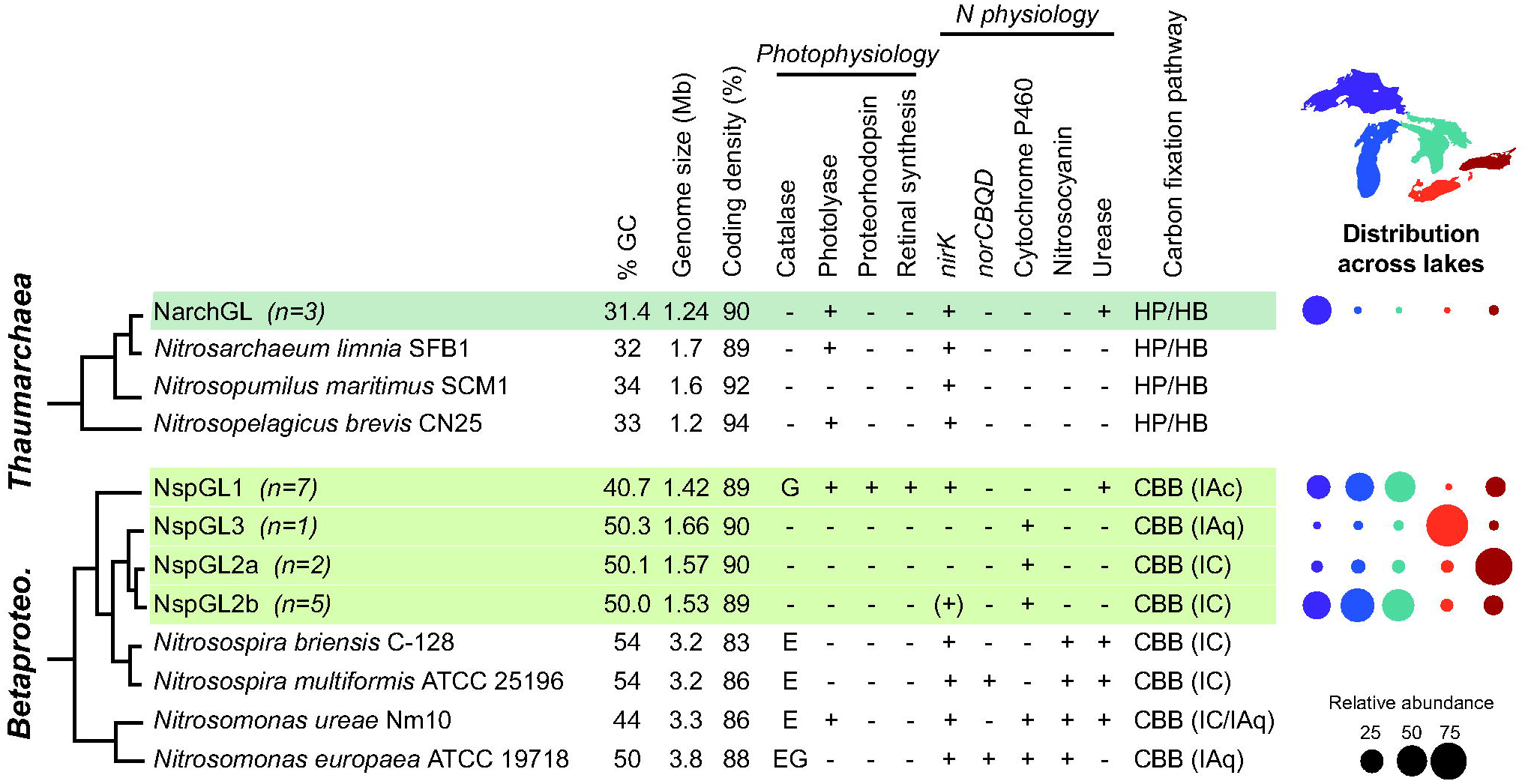
Genome properties and cross-lake distribution of ammonia oxidizing organisms, showing both Archaea (top) and Betaproteobacteria (bottom). Rows highlighted in green represent clusters of genomes reconstructed from the Great Lakes and median values are shown for genome size, GC content, and coding density. For catalase, “E” indicates monofunctional catalase *katE*; “G” indicates bifunctional catalase-peroxidase *katG*. For carbon fixation, RuBisCO type is shown in parentheses (61); HP/HB, 3-Hydroxypropionate/4- Hydroxybutyrate cycle; CBB, Calvin-Benson-Bassham cycle. Bubble plot shows composition of ammonia oxidizers in hypolimnion samples, using MAGs as probes to recruit metagenomic reads (values sum to 100% for each lake column). Genes identified in only a subset of genomes are shown as (+).

**Table 1.**
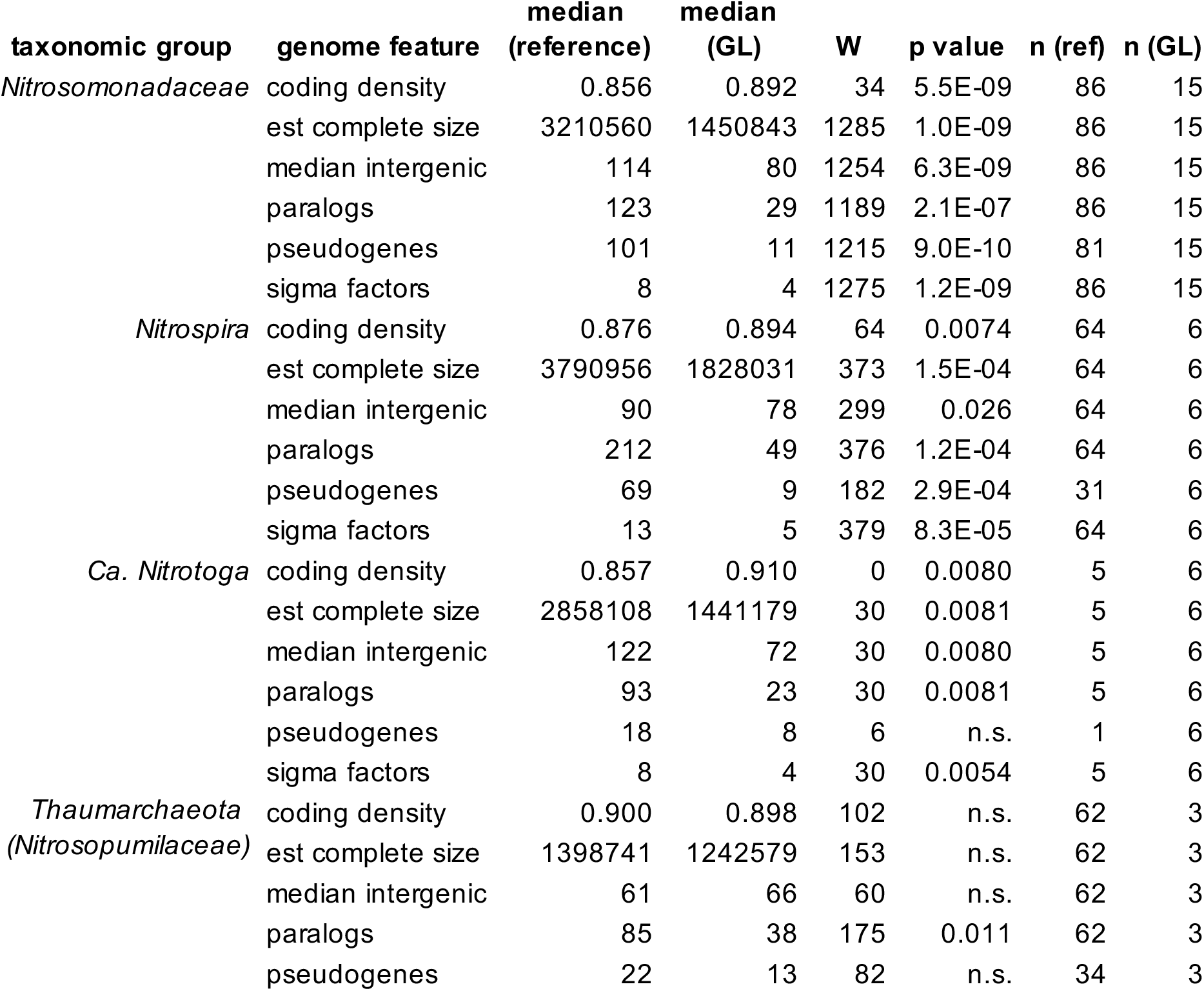
Evidence for genome streamlining in nitrifiers from the Laurentian Great Lakes. Genome features were compared between Great Lakes MAGs and reference genomes using a two-sided Wilcoxon/Mann-Whitney test. n.s., not significant at 0.05 level. Only genomes with >70% completion and <10% contamination are included.

We next compared gene content between our Great Lakes *Nitrosospira* and 86 *Nitrosomonadaceae* reference genomes. On average, Great Lakes *Nitrosospira* encode far fewer two-component signal transduction systems (NspGL = 5-8, mean reference = 19), transposases (NspGL = 0-7, mean reference = 39), motility genes (NspGL = 0-4, mean reference = 52), pilus and secretion genes (NspGL = 2-9, mean reference = 27), and defense-related genes (NspGL = 4-11, mean reference = 39) (Dataset S5). They also lack functions related to biofilm formation such as polysaccharide matrix production (e.g. *pel* genes) and extracellular protein targeting (exosortase and PEP-CTERM motifs). We note that while our 15 new *Nitrosospira* MAGs have high estimated completion (median 98.6%; Dataset S3), some gene absences may reflect incomplete assemblies. Nevertheless, this overall picture of gene content in Great Lakes AOB contrasts with *Nitrosospira* isolates from soil (45, 47) and even to oligotrophic *Nitrosomonas* isolates (49), and is consistent with a passive planktonic lifestyle in extremely low-nutrient systems.

We next compared metabolic potential among Great Lakes AOB ecotypes to understand their ecological preferences for upper lakes (NspGL1, NspGL2b), Ontario (NspGL2a), and Erie (NspGL3). Surprisingly, all seven NspGL1 MAGs encode proteorhodopsin, a light-driven proton pump that supports bacterial energy production (50, 51). They also carry the genes necessary to synthesize its chromophore retinal, including 15,15’-Beta-carotene dioxygenase (*blh*), lycopene cyclase (*crtY*), phytoene dehydrogenase (*crtI*), phytoene synthase (*crtB*), and GGPP synthase (*crtE*) ((52, 53); Fig 3a). We also identified proteorhodopsin in a single-cell amplified genome representing NspGL1 from Lake Michigan (Fig 3a; 99.8% ANI with NspGL1 MAGs), demonstrating that it is not an artifact of metagenome assembly. To our knowledge, this is the first example of a nitrifier with proteorhodopsin. All NspGL1 proteorhodopsins share residues H95, D127 and E138 along with a short beta-turn (G111-P116) between helices B and C, which are characteristic features of proteorhodopsin as distinct from sensory and other rhodopsins (54), and the presence of leucine at position 135 suggests green light tuning (55) (Fig 3b). All of the genes in this module have highest similarity to homologs from *Polynucleobacter*, but are flanked by Nitrosomonadaceae-like genes, suggesting recent horizontal gene transfer (Fig 3a). The predicted NspGL1 proteorhodopsins cluster with *Polynucleobacter*, *Methylopumilus*, and other freshwater Betaproteobacteria in Supercluster III as defined by MicRhoDE ((56); Fig 3c). We compared the homologous genome region in two highly similar MAGs from Lakes Biwa and Baikal (Fig S5); these contigs lack the proteorhodopsin module but appear to flank a variable region where the contig assembly ends. A proteorhodopsin photosystem could support survival of NspGL1 in the presence of sunlight, which has been shown to inhibit ammonia oxidation (37, 57). In the upper lakes where NspGL1 is abundant, light penetration is high well below the thermocline in stratified periods (58), and deep water taxa are seasonally advected to the surface by water column mixing (34). In addition to proteorhodopsin, NspGL1 — but not the other three ecotypes of Great Lakes *Nitrosospira* — encode a class I cyclopyrimidine dimer photolyase, which uses light energy to repair UV-induced DNA damage, and the catalase-peroxidase *katG*, suggesting that the NspGL1 ecotype is adapted to relatively shallow depths in the water column (Fig 2).

**Figure 3.**
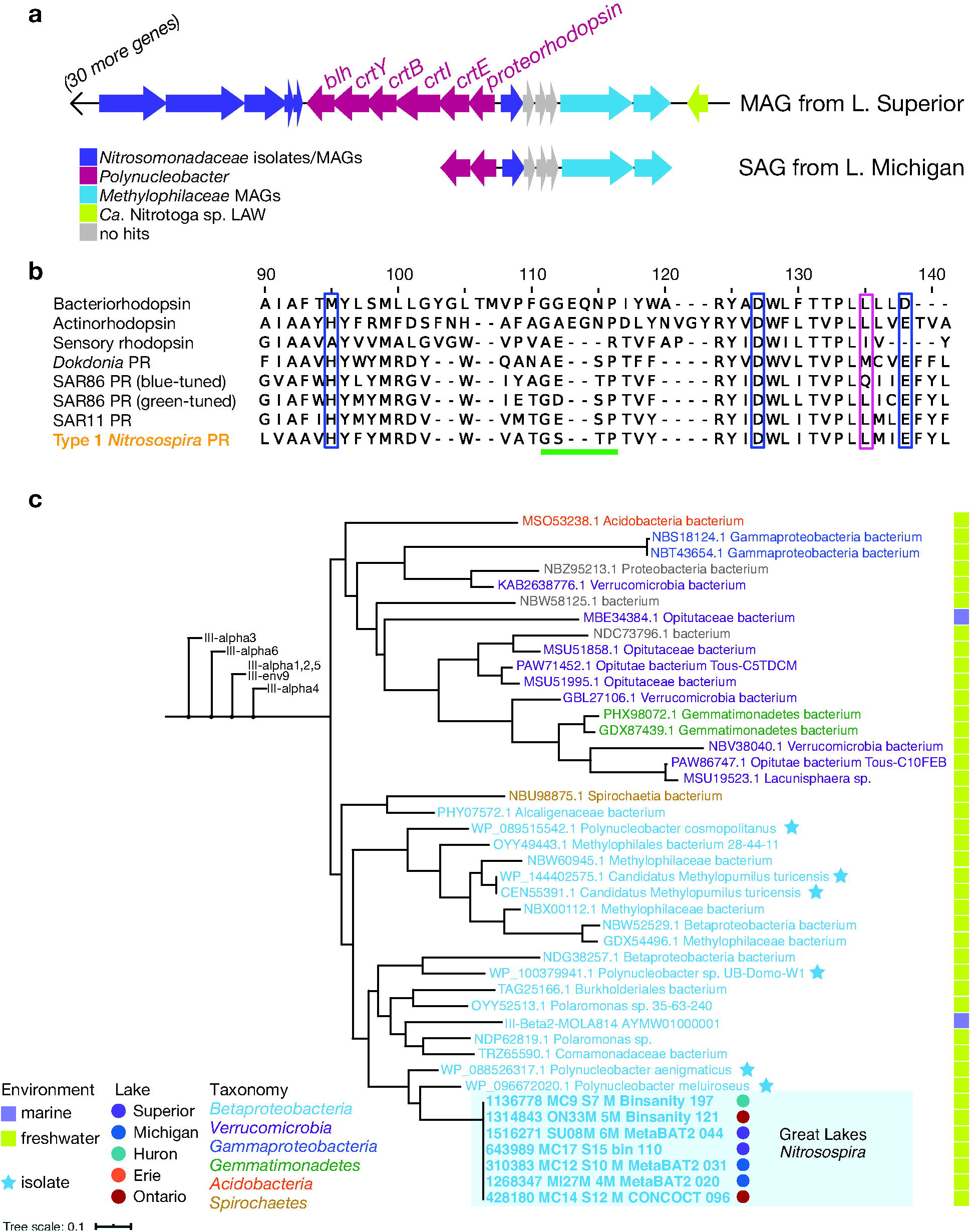
Evidence for proteorhodopsin (PR) in *Nitrosospira* from the Great Lakes. (a) Gene neighborhood surrounding PR in *Nitrosospira* MAG MC17_S15_bin_110 and SAG 207399. Genes are colored according to the best BLAST hit taxonomy in the NCBI nr database. (b) Alignment of predicted *Nitrosospira* PR with reference sequences. Diagnostic features are highlighted (54, 55): blue boxes, diagnostic residues for PR; pink box, residue indicative of blue or green tuning; green underline, shorter beta-sheet region in PR. Sequence accession numbers: bacteriorhodopsin P02945, actinorhodopsin A0A1D9E0H1, sensory rhodopsin P42196, *Dokdonia* PR EAQ40507.1, SAR86 blue-tuned PR Q4PP54, SAR86 green-tuned PR Q9F7P4, SAR11 PR A6YQL7. (c) Phylogenetic tree showing close relatives of *Nitrosospira* PR within Supercluster III, as defined by MicRhoDE database (56). Neighboring clusters have been collapsed for clarity.

Great Lakes *Nitrosospira* carry a reduced, ecotype-specific complement of nitrogen metabolism genes compared to reference AOB (Fig 2, Dataset S5; gene absences were verified as described in Materials & Methods). All are presumed to have the core ammonia oxidation enzymes ammonia monooxygenase and hydroxylamine dehydrogenase; these genes were assembled and binned as expected in some MAGs, and were manually identified on short unbinned contigs in other cases (Dataset S6; see Materials & Methods). Surprisingly, all Great Lakes *Nitrosospira* MAGs lack the copper protein nitrosocyanin, whose precise function is unknown but so far has been found in all described AOB except one member of the *N. oligotropha* clade (49). Based on the expanded set of genomes analyzed here, the lack of nitrosocyanin likely extends beyond the Great Lakes MAGs to closely related freshwater and marine strains, along with five more members of the *N. oligotropha* clade (Fig S3); its absence may be related to the divergence of these clades. Only NspGL1 and NspGL2b encode NO-forming nitrite reductase (NirK), which confers nitrite tolerance (59); this result is surprising given that these two clades dominate the upper lakes where productivity and reduced N are lowest. None of the Great Lakes ecotypes encode NO reductase (NorCBQD) and NspGL1 lacks cytochrome P460 family proteins, both of which are common in AOB and implicated in nitrogen oxide metabolism (18, 49). Nitrogen acquisition is also distinct among Great Lakes AOB: NspGL1 lacks an apparent ammonium transporter, but encodes urease structural and accessory genes (*ureABCEFG*) and a high-affinity urea transporter (*urtABCDE)*. Further, all Great Lakes ecotypes encode a high-affinity amino acid transporter (*livFGHM*); these genes are rare (<5%) in reference genomes and could supply reduced nitrogen and/or organic carbon. Finally, NspGL1 and NspGL3 have genes for producing cyanophycin, an intracellular storage compound for nitrogen (47, 60). Together, the distinctive gene complements present in Great Lakes *Nitrosospira* illustrate the variability and adaptability of AOB gene content, even across a connected freshwater habitat.

As with nitrogen metabolism, carbon metabolism is also distinct between Great Lakes and reference AOB, and among Great Lakes ecotypes (Fig 2, Dataset S5). Unlike most reference AOB, Great Lakes *Nitrosospira* lack two key enzymes of the oxidative pentose phosphate pathway, glucose-6-phosphate dehydrogenase and 6-phosphogluconate dehydrogenase. All ecotypes except Erie-specific NspGL3 also lack genes for glycogen synthesis and degradation, suggesting that they are unable to store and access this carbon reserve. The key enzyme for carbon fixation, RuBisCO, has evolved several kinetically distinct forms whose distribution likely reflects ecological pressures (61). NspGL1 and NspGL3 both contain Form IA RuBisCO, while NspGL2a and NspGL2b contain Form IC RuBisCO (Fig 2; (61, 62)). NspGL1 genomes also possess an alpha carboxysome-like *cso* operon, similar to *Nitrosomonas eutropha* C91 (62), though our draft assembly lacks the expected carbonic anhydrase (*csoS3*/*csoSCA*). Carboxysome-associated RuBisCO may allow NspGL1, the ecotype most strongly adapted to energy and nutrient limitation, to more efficiently fix CO_2_ by minimizing the wasteful oxygenation reaction and reducing the cellular nitrogen allocation to RuBisCO (61). The ranges of kinetic properties observed in other autotrophs for Form IAq (found in NspGL3) and Form IC (found in NspGL2a and NspGL2b) overlap, and therefore more work is needed to understand the fitness advantages, if any, that this RuBisCO diversity confers on Great Lakes nitrifiers.

### Streamlined freshwater Thaumarchaeota

We reconstructed three similar genomes (>99% ANI) of *Nitrosarchaeum* (NarchGL; Fig S6, Dataset S3) from three separate samples (two from Superior, one from Ontario), consistent with our low observed 16S rRNA diversity for *Thaumarchaeota*. These NarchGL genomes are very similar (∼99% ANI) to two genomes reconstructed from Lake Baikal, located thousands of kilometers away (63). Their next closest relatives are also from freshwater environments, and phylogenetic clustering suggests that salinity is an important driver of divergence throughout the *Nitrosopumilaceae* (Fig S6). As a group, the *Thaumarchaeota* tend to have smaller genomes, lower G+C content, higher coding density, and fewer paralogs and pseudogenes than nitrifying bacterial taxa; even within this group, NarchGL genomes fall below the 30th percentile in size and have significantly fewer paralogs than average (Table1, Fig S4, Dataset S4). Using our reconstructed genomes as probes for metagenomic read recruitment, NarchGL were detected in Superior, Michigan, and Ontario; they represented roughly one-third of ammonia oxidizers in the mid-hypolimnion of station SU08M (Fig 2).

NarchGL share nearly 90% of their predicted proteins with close relatives including *Ca.* Nitrosarchaeum limnia; at the same time, they show distinctive patterns in gene content that pinpoint the key selective pressures of deep lakes (Fig 2, Dataset S5). All three NarchGL genomes encode urease and a urea transporter, implicating urea as a vital source of nitrogen for energy and/or biosynthesis. Consistent with phosphorus scarcity in much of the Great Lakes (64), NarchGL encode high affinity transport systems for phosphate and potentially phosphonates, though we did not identify a phosphonate lyase. In addition to both CRISPR/Cas enzymes cas1 and cas4, NarchGL genomes contain several phage proteins, suggesting viral infection and integration events may be common. DNA photolyases, which have been found in epipelagic clades of marine *Thaumarchaeota* (7), are present in all low salinity *Nitrosarchaeum* including NarchGL, suggesting NarchGL are exposed to sunlight due to high water clarity (58) and/or annual mixing in the Great Lakes. NarchGL also lack the common tRNA modification 4- thiouridylation (indicated by K04487 and PF02568-PF18297 (65)); we propose that the absence of this modification, which is susceptible to near-UV radiation (65), is also related to sunlight exposure.

Genomes of NarchGL reveal striking reduction in environmental sensing, response, and regulatory functions, relative to most other *Nitrosarchaeum* and *Nitrosopumilaceae* (Dataset S5). NarchGL encode 9-12 domains representing the general archaeal transcription factors, TATA binding protein (TBPs, PF00352) and transcription factor B (TFBs, PF00382 and PF08271), compared to 21 in *Ca.* N. limnia. NarchGL lack common domains found in two-component systems that transmit environmental signals to control gene expression or protein activity (domains PF02743, PF00672, PF00512, PF00072; NarchGL = 0, *Ca.* N. limnia = 53-54 copies per genome). Further, they are depleted in ArsR family transcription factors (PF01022; 0 copies in NarchGL vs. 2-3 in *Ca.* N. limnia), P-II proteins for regulation of nitrogen metabolism (PF00543; 1 copy per NarchGL genome vs. 5 in *Ca.* N. limnia), and other potential regulatory domains (CBS PF00571: 5 in NarchGL vs. 18-19 in *Ca.* N. limnia; USP PF00582: 1 in NarchGL vs. 15 in *Ca.* N. limnia). This extremely limited regulatory capacity in NarchGL stands in sharp contrast to closely related *Ca.* N. limnia, but instead parallels the oceanic minimalist *Ca.* Nitrosopelagicus brevis (9).

### Expanded diversity of Ca. *Nitrotoga* with reduced genomes

Despite the broad distribution of *Ca.* Nitrotoga in freshwater systems and beyond, only six genomes are available, derived from rivers heavily impacted by urban and agricultural influence, a wastewater treatment plant, and coastal sediment (27, 66, 67). Hence the metabolic and phylogenetic diversity of this group is virtually unexplored. We reconstructed six new MAGs of *Ca*. Nitrotoga, which form two clusters with >99% ANI within each cluster and ∼97% ANI between clusters (NtogaGL1a and NtogaGL1b; Fig S3). These new *Ca*. Nitrotoga MAGs are far smaller than published genomes (median GL = 1.44 Mb, reference = 2.61-2.98 Mb), have shorter intergenic regions, fewer sigma factors, and fewer paralogs (Table 1, Fig S4, Dataset S4), consistent with genome streamlining (48). They lack functions such as motility and chemotaxis, pilus biogenesis, and DNA repair (*mutLS*) (Dataset S5). Great Lakes *Ca.* Nitrotoga also encode markedly fewer two-component systems for sensing and responding to environmental cues than four river-derived genomes (NtogaGL = 2-6 per genome vs. 30-35 in four reference genomes; reference strain KNB has 7). Compared to reference genomes, NtogaGL have fewer defense-related genes (restriction-modification, toxin-antitoxin, and CRISPR-Cas systems; mean NtogaGL = 11 vs. 39 for references), and transposases (mean NtogaGL = 3 vs. 19 for references) (Dataset S5). While incomplete assembly of hypervariable genome regions may explain some of these absences, the overall genome properties are consistent with a relatively stable low-nutrient environment and planktonic lifestyle.

The reduced genomes of NtogaGL1a/b help clarify core features of the genus *Nitrotoga*, along with accessory functions that may enable local adaptation in specific populations. To date, sequenced *Nitrotoga* including NtogaGL1a/b encode similar electron transport pathways, including NADH dehydrogenase-Complex I, succinate dehydrogenase-Complex II, and Alternative Complex III, along with high-affinity cbb3-type cytochrome oxidases suggesting adaptation to low oxygen conditions. They also share the Calvin cycle for carbon fixation, a complete TCA cycle, and an evolutionarily distinct nitrite oxidoreductase (NXR) from other NOB (27, 66, 67). All *Nitrotoga* to date also share transporters for amino acids and peptides, potential sources of C and/or N. *Nitrotoga* can also potentially access reduced sulfur compounds for energy via sulfite dehydrogenase, suggesting metabolic flexibility beyond nitrite oxidation.

Beyond these similarities, the small genomes of NtogaGL1a/b are distinct from previously described *Nitrotoga* in many ways. NtogaGL1a/b lack NiFe hydrogenase to use hydrogen as an energy source. They also lack nitrogen metabolism functions including assimilatory nitrite reductase (*nirBD*) and nitrite reductase to NO (*nirK*). Based on gene content, NtogaGL1a/b appear unable to use hexoses like glucose, since they lack the glycolytic enzyme phosphofructokinase and the Entner-Doudoroff pathway, similar to *Nitrobacter winogradskyi* (68). Consistent with this, they also lack genes for storage and breakdown of glycogen (Dataset S5). All but one of the NtogaGL1a/1b genomes encode cyanate lyase (*cynS*), which is found in other NOB but no *Nitrotoga* to date (25, 69, 70). The *cynS* gene, adjacent to *glnK*-*amtB* for ammonium sensing and transport, likely functions in N assimilation, as recently described for *Nitrospinae* (71). While cyanase has been shown to mediate reciprocal feeding between some NOB and ammonia oxidizers (25), it remains to be seen whether such an interaction occurs in the free-living (<1.6um) size-fraction and dilute environment sampled here. Notably, cyanase from NtogaGL1a/1b, along with predicted *Nitrospirae* proteins from Lake Baikal and soil, form a distinct phylogenetic cluster from most nitrifier cyanase proteins observed to date (Fig S7).

The two ANI-based clusters we detected, NtogaGL1a and NtogaGL1b, appear to be phylogenetically and ecologically distinct ecotypes. Based on short-read mapping, NtogaGL1b dominates Erie, while NtogaGL1a dominates all other *Ca.* Nitrotoga-containing samples (Fig 4). We found several metabolic genes that differentiate the two ecotypes. ER-specific NtogaGL1b genomes share a region encoding thiosulfate dehydrogenase (*tsdA*), cytochromes, transport of sulfur-containing compounds, lactate dehydrogenase (ldh), a two-component system, and a Crp-family transcription factor (Fig S8). This region may be involved in oxidizing thiosulfate as an energy source, and sensing and responding to redox changes that accompany seasonal hypoxia in Lake Erie. The corresponding region in NtogaGL1a encodes an integrase and photolyase, consistent with greater DNA photodamage in the more transparent waters of Michigan, Huron, and Ontario where NtogaGL1a is abundant.

**Figure 4.**
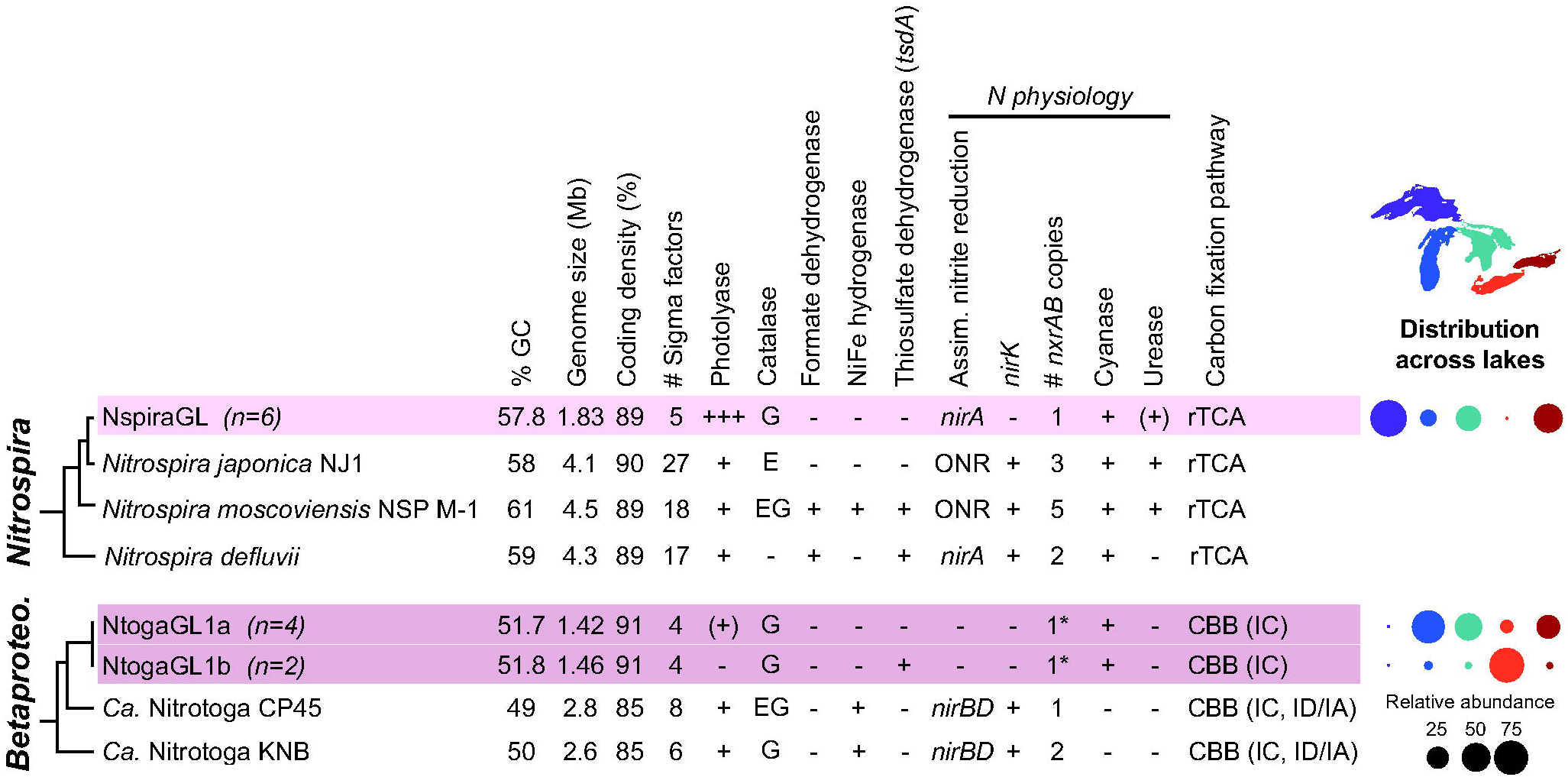
Genome properties and cross-lake distribution of nitrite oxidizing taxa *Nitrospira* (top) and *Ca.* Nitrotoga (Betaproteobacteria; bottom). Rows highlighted in pink represent clusters of genomes reconstructed from the Great Lakes and median values are shown for genome size, GC content, and coding density. rTCA, reductive tricarboxylic acid cycle; CBB, Calvin-Benson-Bassham cycle; ONR, octaheme nitrite reductase. Values in parentheses indicate RuBisCO type (61). Bubble plot shows composition of NOB per lake based on metagenomic read mapping. Genes identified in only a subset of genomes are indicated by (+). The asterisk (*) indicates for *Ca.* Nitrotoga, one *nxrAB* copy was recovered in genome assemblies, but short read analysis suggests two copies per genome (Supplemental Text).

### Great Lakes *Nitrospira* reveal adaptations to sunlit oxic environment

We reconstructed six closely related genomes of *Nitrospira* (∼99% ANI; Fig S9, Dataset S3), representing the predominant NOB throughout Lake Superior and in parts of Michigan, Ontario, and Huron (Fig. 4; Dataset S1). These genomes, which we refer to as NspiraGL, fall within lineage II (Fig S9), which is broadly distributed across soil, freshwater, and engineered habitats (20); however, genome analyses to date have focused on strains from wastewater and engineered systems, leaving major blind spots. NspiraGL share core features of *Nitrospira* metabolism, including a periplasmic-facing NXR that is advantageous under substrate-limiting conditions, multiple cytochrome *bd*-like oxidases, and the reverse TCA cycle for carbon fixation (69). However, as with *Ca.* Nitrotoga, the *Nitrospira* genomes we reconstructed in the Great Lakes are markedly smaller than published reference genomes (median NspiraGL = 1.83 Mb, median reference = 3.72 Mb), with higher coding density and fewer paralogs, sigma factors and pseudogenes (Fig. S4; Dataset S4), consistent with genome streamlining theory (48). Compared to 75 lineage II *Nitrospira* reference genomes, NspiraGL have reduced capacity for environmental sensing (two-component systems: NspiraGL = 7, mean reference = 26), transport (NspiraGL = 76-83, mean reference = 140), defense (NspiraGL = 7-8, mean reference = 26), and transposition (NspiraGL = 0-2, mean reference = 15), and lack pilus or flagellar motility (Dataset S5). NspiraGL encode just five sigma factors, compared to 18 in *N. moscoviensis.* Further, NspiraGL genomes encode a single NXR, while *N. moscoviensis* carries five copies that are differentially regulated (26, 72). NspiraGL also lack the *glnE* gene for glutamine synthetase (GS) adenylyltransferase, suggesting that GS activity is not repressed by this mechanism. Together, these features suggest limited regulatory and ecological flexibility, consistent with a relatively constant, oligotrophic environment.

Compared to other *Nitrospira,* NspiraGL exhibit limited energetic flexibility, but can access diverse nitrogen sources (Fig. 4, Dataset S5). We predict that NspiraGL are unable to grow on hydrogen or formate as alternative energy sources (23, 26), as they lack NiFe- hydogenase and formate dehydrogenase. The glycolysis and oxidative TCA cycles appear to be incomplete, lacking phosphofructokinase and citrate synthase, respectively; this suggests a limited capacity for organic carbon utilization. NspiraGL lack *nirK,* encoding NO-forming nitrite reductase, which is found in a majority of reference genomes. To obtain nitrogen for biosynthesis, NspiraGL encode a high-affinity nitrate/nitrite/cyanate transporter (*nrtABC*), assimilatory nitrite reductase (*nirA*), and cyanase (*cynS*), along with *amt* family ammonium transporter. Although none of the NspiraGL MAGs include urease (*ureCBA*), one does contain urease accessory proteins (*ureEFGD*) and two contain a urea transporter (*urtABCD*), suggesting incomplete assembly of the urea utilization pathway. As with *Ca.* Nitrotoga, we suggest that cyanase, along with urease where present, functions in nitrogen assimilation rather than cross-feeding, given the dilute environment and free-living planktonic cells.

Beyond energy, carbon, and nitrogen metabolism, we discovered striking differences between NspiraGL and reference *Nitrospira* related to DNA repair. NspiraGL encode two additional photolyase-related proteins, along with a class I cyclopyrimidine dimer (CPD) photolyase found in most reference *Nitrospira* (Fig 5). Photolyases use blue light energy to repair DNA lesions caused by UV radiation (73). The two additional genes in NspiraGL are adjacent and share best hits with *Betaproteobacteria*, suggesting recent horizontal transfer (Fig S10). One likely encodes an FeS-BCP photolyase, which repairs (6–4) dipyrimidine lesions (74, 75). The other shares an FAD-binding domain with photolyases but the C-terminal region has no recognizable domains (Fig 5). This protein is widespread in aquatic bacteria and has not been functionally characterized, though an actinobacterial homolog was suggested to be involved in light sensing and regulation (76). Beyond photolyases, NspiraGL also encode uracil-DNA glycosylase (UNG), which removes misincorporated uracil from DNA. Uracil results from deamination of cytosine, which can occur spontaneously or be induced by NO (77). In addition to the photolyases and UNG that repair DNA lesions, NspiraGL encode translesion DNA polymerase V (*umuCD*) which enables replication to proceed past lesions. Together, these genes indicate that *Nitrospira* in the Great Lakes experience significant DNA damage, including UV-induced damage that also requires light for the repair process, in hypolimnion waters with high transparency (58) and/or during seasonal mixing.

**Figure 5.**
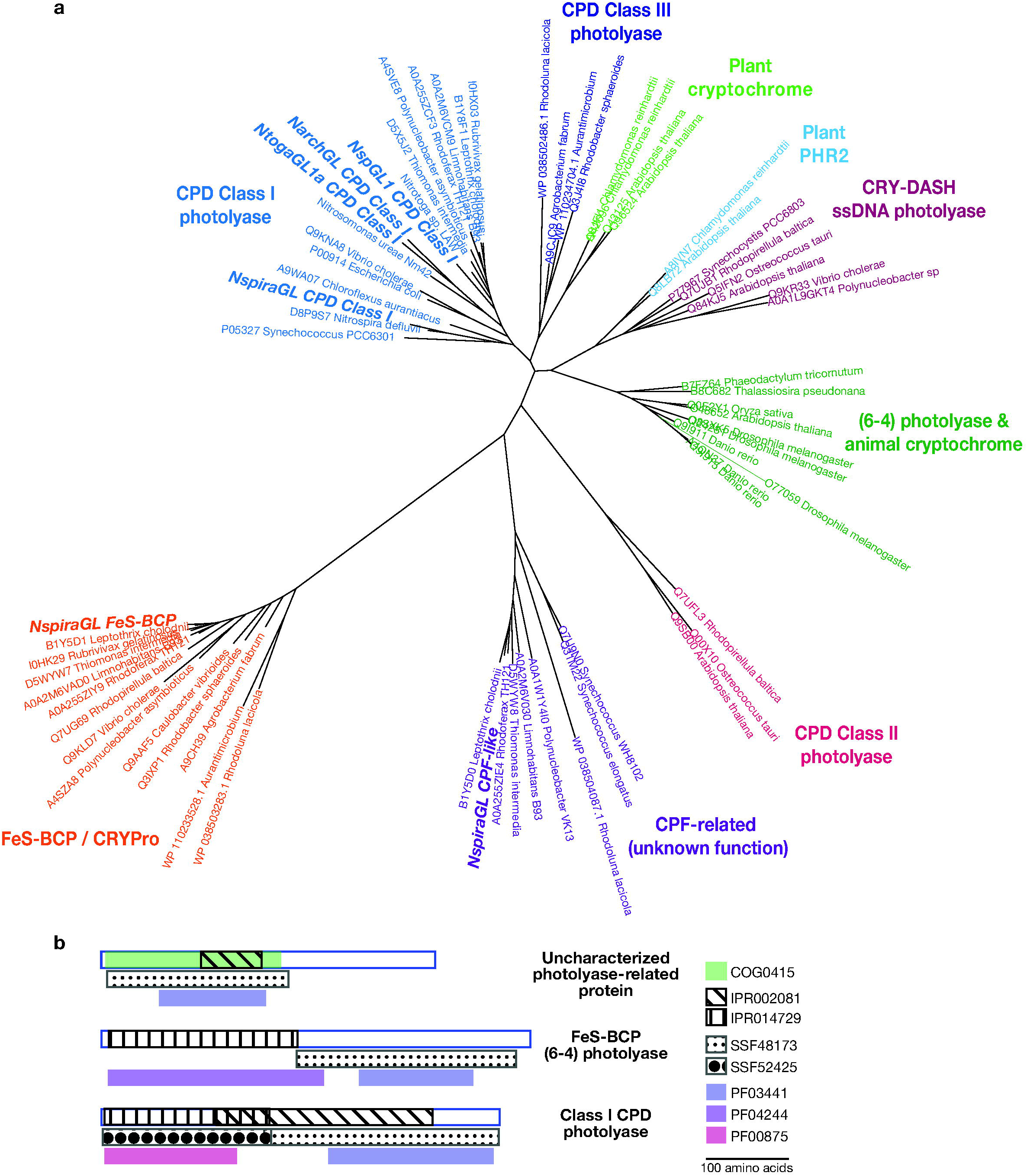
Distinct photolyase proteins in NspiraGL. **a)** Phylogenetic tree showing families of photolyases. Three families are found in NspiraGL: CPD class I photolyase, FeS-BCP / CRYPro family, and an uncharacterized CPF-related family found in diverse Bacteria. CPD Class I photolyases are also found in other nitrifiers including *Ca*. Nitrotoga NtogaGL1a, *Nitrosospira* NspGL1, and *Nitrosarchaeum* NarchGL. **b)** Domain structure of the three photolyase families present in NspiraGL.

Other major differences between NspiraGL and reference *Nitrospira* are related to reactive oxygen species (ROS) (Dataset S5). Surprisingly, despite their oxic habitat, NspiraGL lack superoxide dismutase (SOD), monofunctional catalase (*katE*), and bacterioferritin, which limits the Fenton reaction by sequestering free iron. However, all six NspiraGL MAGs, but few reference genomes (7% of 75), have recently acquired bifunctional catalase-peroxidase *katG*; interestingly we also observed *katG* in Great Lakes *Ca.* Nitrotoga and *Nitrosospira* (Fig 2, Fig 4). The absence of SOD suggests that NspiraGL does not produce damaging levels of endogenous superoxide, perhaps because NspiraGL lack the major respiratory and non-respiratory flavoproteins that produce ROS in other SOD-containing *Nitrospira* (78). Unlike superoxide, H_2_O_2_ can cross membranes, and is known to be produced by both photooxidation of dissolved organic matter and dark heterotrophic activity (79). The lakes where NspiraGL dominate have high water clarity (58) and low productivity and are fully oxic, consistent with abiotic photochemistry as the primary source of exogenous ROS; this stress may have selected for *katG* as a defense. NspiraGL also lack cytochrome *c* peroxidase, which is found in 70 of 75 reference genomes; this protein is proposed to function in anaerobic respiration of H_2_O_2_ (80) and therefore its absence in NspiraGL is consistent with a constant oxic environment. Together, these results indicate that *Nitrospira* in the Great Lakes face distinct ROS pressures that have shaped their gene content.

## Conclusions

The Laurentian Great Lakes harbor nitrifiers that are phylogenetically related, but markedly different in genome size and functional capacity, from their well-studied relatives inhabiting wastewater systems, soils, and even other freshwater systems. By examining the entire nitrifier assemblage at once, we detected common features across taxa that illuminate the selective pressures faced by microbes in deep lakes. All the lineages we describe show small genome sizes (1.3-1.7 Mb), reduced capacity for environmental sensing and response, and adaptation to a passive (i.e. non-motile) planktonic lifestyle, features which have not been previously associated with AOB, *Nitrospira*, and *Ca.* Nitrotoga. Within the AOB *Nitrosospira*, we found ecotypes with a gradient of genome reduction that maps onto their habitats’ trophic gradient: from NspGL1 (1.4 Mb, low GC, upper lakes) to NspGL2b (1.5 Mb, upper lakes) to NspGL2a (1.6 Mb, Ontario) to NspGL3 (1.7 Mb, Erie) (Fig 2). The thaumarchaeal NarchGL have markedly reduced regulatory capacity like the open ocean strain *Nitrosopelagicus brevis* (9). The NOB NspiraGL have genomes 50-60% smaller than described *Nitrospira* and dominate, along with the AOB NspGL1 and the AOA NarchGL, the deeper more oligotrophic basins, while *Ca.* Nitrotoga favor shallower, more productive basins. The emergence of Erie-specific ecotypes of both *Nitrosospira* (NspGL3) and *Ca.* Nitrotoga (NtogaGL1b) demonstrates how distinct this habitat is compared to the other lakes. Importantly, our findings here represent planktonic cells in the smallest size fraction (<1.6 µm); it is likely, especially in Erie, that particle-associated nitrifiers may be abundant and genetically distinct.

Nitrifiers inhabiting the transparent waters of the upper Great Lakes show distinctive adaptations to light including diverse photolyases, ROS detoxification, and even proteorhodopsin. This discovery is surprising, given that nitrifiers are rare in the surface mixed layer of the Great Lakes (Fig. S1) and that photoinhibition of ammonia oxidation and nitrifier growth is well documented (37, 40, 57). We propose that proteorhodopsin could be used to augment energy metabolism when ammonia oxidation is photoinhibited and/or ammonia is substrate limited. Water clarity has increased over the past several decades in Lakes Michigan and Huron, now surpassing that of Lake Superior (58). High light penetration along with seasonal mixing likely exposes deep water cells to damaging levels of light and oxidative stress. Future cultivation and physiological studies should examine photoinhibition and potential phototrophy in Great Lakes nitriifiers.

The capacity for nitrification is found across multiple phyla, and our work unveils new clues to understanding the ecological and evolutionary potential of these diverse lineages. This collective nitrifier diversity undoubtedly influences the cycling of carbon and nitrogen across this ecosystem, and future work will explore the differential contributions to nitrification by the distinct lineages we described here. Understanding what controls the diversity of nitrifiers and other key functional groups, and the consequences of this diversity for biogeochemistry, are essential for forecasting the effects of rapid environmental change across the large lakes of the world (e.g.(81)) and predicting impacts on the critical ecosystem services they provide (82).

## Materials & Methods

### Sample collection

Water samples were collected from the Laurentian Great Lakes aboard the R/V *Lake Guardian*, during the biannual Water Quality Surveys conducted by the U.S. EPA Great Lakes National Program Office (83). Station information is provided in Supplemental Dataset S8. Data presented here were collected in April and August 2012. Samples were collected using a CTD rosette sampler (Sea-Bird Scientific) at the surface (2 m), deep chlorophyll maximum (if present), the mid-hypolimnion (depths ranging from 19 m in Erie to 200 m in Superior; Dataset S1), and near the bottom of the water column (10 m above the lake bottom at most stations, 1 m above bottom at shallow stations). For each sample, 5-8L of water was pre-filtered through a GF/A glass fiber filter (Whatman 1820-047; nominal pore size 1.6 μm) to exclude eukaryotic phytoplankton and particle associated microbes, and cells were collected on 0.22 μm Sterivex filters (Millipore SVGP01050). Filters were stored at -80°C. For dissolved nutrient analysis, 0.22 μm filtrate was collected in 125 ml acid-clean HDPE bottles (Nalgene) and stored at -20°C. Samples for single cell amplified genomes (SAG) were collected in August 2014. For each sample, 1 ml of raw water was incubated with 100 µl of glycerol-TE buffer (20 ml 100X TE pH 8 + 100 ml glycerol + 60 ml water; final concentration after sample addition is 10mM Tris, 1mM EDTA, 5% glycerol) for 10 minutes in the dark, then flash frozen in liquid nitrogen and stored at - 80°C until processing.

### Physicochemical data

CTD profiles, water chemistry and chlorophyll-*a* data were collected by the U.S. EPA according to standard protocols (84) and retrieved from the Great Lakes Environmental Database (https://cdx.epa.gov/) for 2012 and 2013. In addition, we measured dissolved nitrogen species from August 2013 samples. Ammonium concentrations were measured using the OPA method in a 96-well plate (85). Nitrate and nitrite concentrations were measured using the Greiss reaction method in a 96-well plate (86). Urea concentrations were measured in a 24 well plate using a colorimetric reaction (87).

### 16S rRNA analysis

The full 16S rRNA amplicon dataset was described by Paver and colleagues (34). Here we focus on data from the V4-V5 region (primers: 515F-Y, 926R (88)), collected in 2012 in tandem with metagenome samples from select stations. We classified sequences using the Silva v. 132 database (89) and the wang method (90) as implemented by mothur (91). Sequences classified to each detected family of nitrifiers (ammonia oxidizer families *Nitrosomonadaceae* and *Nitrosopumilaceae*; nitrite oxidizer families *Gallionellaceae* and *Nitrospiraceae*) with a mothur-assigned confidence score above 90 were delineated into taxonomic units using minimum entropy decomposition with a minimum substantive abundance of 10 (92).

### Metagenome and single-cell genome sequencing

One station per lake in Superior, Michigan, Huron, and Ontario, and two stations in Erie, were selected for metagenome sequencing. Spring 2012 metagenome samples were collected from the surface, and Summer 2012 metagenome samples were collected from the mid- hypolimnion (depths listed in Dataset S1). DNA was extracted using a modified phenol:chloroform extraction protocol (34) and libraries prepared according to the Illumina TruSeq protocol. Samples from spring 2012 were sequenced at the Joint Genome Institute using Illumina HiSeq (2×150bp). Samples from summer 2012 were sequenced at the University of Chicago Functional Genomics Core Facility using Illumina HiSeq 2500 (2×250bp).

To confirm the presence of proteorhodopsin, we analyzed a single cell amplified genome from *Nitrosospira* collected from Lake Michigan and sequenced by the Joint Genome Institute. Quality filtered reads were downloaded from JGI IMG/ER and normalized using bbnorm.sh with target=100 and mindepth=2. Normalized reads were assembled using SPAdes 3.1.11 in single cell mode (93) with flags --sc and --careful. Resulting scaffolds were annotated identically to MAGs as described below.

### Obtaining metagenome-assembled genomes

Raw reads for spring surface samples were quality controlled at the Joint Genome Institute, using bbduk.sh for adapter trimming (ktrim=r, minlen=40, minlenfraction=0.6,mink=11, tbo, tpe, k=23, hdist=1, hdist2=1, ftm=5) and quality filtering (maq=8, maxns=1, minlen=40, minlenfraction=0.6, k=27, hdist=1, trimq=12, qtrim=rl). Raw reads for summer hypolimnion samples were adapter trimmed, quality filtered, and interleaved using bbduk (parameters: ktrim=r, mink=8, hdist=2, k=21, forcetrimleft=10, forcetrimright=199, minlen=150) using BBTools suite version 35.74 (https://sourceforge.net/projects/bbmap/). Separate assemblies of quality filtered reads were carried out for each metagenome using metaSPAdes 3.1.11 --meta mode using default k sizes of 21, 33, 55 (94). To enable binning based on sequence coverage, forward and reverse reads were merged using bbmerge in BBtools, using qtrim2=r trimq=10,13,16 and adapter=default. Merged short reads were then mapped onto each assembly using bowtie2 2.2.9 in --sensitive mode (95), and this coverage information was used to bin assembled contigs. Binning was performed using MetaBAT2 2.12.1 (96), Binsanity 0.2.6.3 (97) and CONCOCT 1.0.0 (98) using default parameters. The resulting bins were scored, aggregated, and de-replicated using DAS_Tool 1.1.1 (99) followed by manual curation using Anvi’o 4.0 (100). We assessed genome completion and contamination of manually curated bins using CheckM 1.1.0 lineage_wf (101), and all new MAGs presented here are greater than 70% complete with less than 10% contamination (Dataset S3). Potential nitrifiers were screened by searching for ammonia monooxygenase, hydroxylamine oxidoreductase and nitrite oxidoreductase within reconstructed genomes using blastp 2.5.0 (102). For bins where any of these genes were detected, we identified bacterial single copy core genes (103) or archaeal single copy core genes (104) using HMMER (105), as implemented in Anvi’o. Single copy core genes were queried against proteins predicted from bacterial and archaeal genomes in RefSeq (NCBI) (106), and taxonomic identity of these core genes was ascertained based on a least common ancestor approach using a 0.1% window around the bit score of the best hit using KronaTools 2.7.1 (107). Taxonomic assignment was further validated using GTDB-tk 1.0.0 (108). Grouping of MAGs into clades and subclades based on ANI was carried out using fastANI 1.1.0 (109). Genome characteristics for each genome group were calculated as the median of those values for the group. Estimated complete genome size was calculated for MAGs and for references in the pangenome analysis using CheckM (101) completion and contamination, as follows: Estimated = Actual*(1-Contamination) / Completion. To quantify the abundance of each clade/ecotype across samples, we used competitive mapping of merged short reads using bowtie2 in sensitive mode against all nitrifier MAGs, summing up mapped read count across all MAGs in a given clade/ecotype, and dividing by total mapped nitrifier reads in a sample; these values are shown in bubble plots (Fig 2, 4).

### Annotation and gene cluster analysis

Reference genomes were obtained from GenBank (accession numbers listed in Supplemental Dataset S4). The full pangenome analyses included all the genomes listed in Supplemental Dataset S4, but we only report results from the subset of genomes most closely related to our MAGs. This subset consists of 86 *Nitrosomonadaceae*, 5 *Ca.* Nitrotoga, 78 *Nitrosopumilaceae* within Thaumarchaeota, and 75 *Nitrospira* that fall within Lineage II. Reference genomes were treated consistently with GL MAGs, with *de novo* gene calling by prodigal 2.6.3 (110) via Anvi’o. Unless otherwise noted, default settings were used for all software. Genes were annotated using InterProScan 5.30-69.0 (111), GhostKOALA (112) and eggnog-mapper 1.0.3 against the bactNOG database (113). Gene cluster analysis was carried out using the Anvi’o pangenome pipeline (114), using blastp to determine sequence similarity, ITEP to eliminate weak similarity (115) and MCL to cluster, using a minbit of 0.5, MCL inflation of 2 and minimum gene occurrence of 1 (116). Sigma factors were tallied by identifying gene clusters annotated with the following PFAMs: PF00309, PF03979, PF00140, PF04542, PF04539, PF04545, PF08281. Pseudogene counts were retrieved where available from NCBI PGAP annotated genomes (117). Paralog counts are reported as the number of gene clusters with more than one gene per genome. Intergenic spacers were calculated using bedtools complementBed function (118). Prokka 1.14.5 (119) was used to generate GenBank-format files from MAGs and SAGs, and genoPlotR 0.8.9 (120) was used to generate initial gene neighborhood maps.

### Gene tree construction

The NspGL1 proteorhodopsin sequence was inserted into the MicRhoDE rhodopsin tree using pplacer (121) through the MicRhoDE Galaxy pipeline (56). We then constructed a more targeted phylogenetic tree using aligned reference sequences of Supercluster III from MicRhoDE, filtered to exclude fragments shorter than 220 amino acids. To this alignment, we added NspGL1 sequences using MAFFT 7.310 (122) along with high similarity sequences from NCBI nr that were not present in MicRhoDE. The tree was inferred using RaxML 8.2.12 with model PROTGAMMALG (123). The tree was visualized in iTOL (124) and more distant clusters were collapsed for clarity.

A cyanase phylogenetic tree was created using sequences drawn from querying NtogaGL cyanase against NCBI nr using blastp, as well as sequences from references (2, 25, 125). Sequences were aligned using MAFFT (122) and the tree was inferred using RaxML 8.2.12 with model PROTGAMMALG (123). Tree was visualized in iTOL (124) and branches were colored based on the taxonomy of the parent genome.

Photolyase-related proteins in GL MAGs were identified by searching for the following features: K01669, COG0415, PF03441, PF00875, PF04244, SSF48173, SSF52425. Reference proteins (n=56) spanning the previously defined families of photolyases and cryptochromes (126) were obtained from UniProt, along with aquatic bacterial sequences described by Maresca and colleagues (76). The reference sequences were aligned using MAFFT (122), and sequences from GL MAGs were added using the MAFFT --addfragments option. The tree was estimated using IQ-TREE 2 1.6.11 (127) and visualized using iTOL (124).

### Phylogenomic tree construction

*Nitrospirae*, *Thaumarchaeota*, *Gallionellaceae* and *Nitrosomonadaceae* genomes were downloaded from Genbank (NCBI) (128) and included in phylogenomic trees for their respective family. Phylogenomic analyses were carried out within Anvi’o. Briefly, single copy core genes were extracted as described above, individually aligned at the protein level using muscle (129), and concatenated for each genome. Concatenated alignments were trimmed using Gblocks 0.91b (130) and analyzed by RAxML 8.2.12 (123) to create a phylogenetic tree using the PROTGAMMALG model and 50 bootstraps. Trees were visualized in iTOL (124).

### Proteorhodopsin assembly verification

We used several approaches to validate the presence of proteorhodopsin in assembled *Nitrosospira* genomes, to rule out the possibility of chimeric assemblies from different species. We note that proteorhodopsin-containing contigs were independently assembled and binned together with core *Nitrosospira* contigs from seven different samples (i.e. each sample was assembled and binned separately, rather than co-assembled). In five of seven cases, proteorhodopsin and retinal biosynthesis genes were assembled together with core *Nitrosospira* genes on the same contig. To rule out a systematic reproducible error in assembly and/or binning, we compared these seven MAGs to a single cell amplified *Nitrosospira* genome (SAG) from Lake Michigan, obtained as part of another project with the JGI. This SAG was processed through JGI’s standard decontamination pipeline and manually investigated to ensure lack of contamination. We found no evidence of contaminating core genes, as all core genes had best hits to either *Nitrosospira* or more generally *Nitrosomonadaceae* in nr. SAG contigs were matched to homologous contigs from NspGL1 MAGs to determine if any SAG contigs were unique using FastANI 1.1.0 (109) with --visualize flag. All contigs from this *Nitrosospira* SAG were found within an NspGL1 MAG. Bandage 0.8.1 (131) was used to manually inspect the assembly graph around the contig that contained the NspGL1 *Nitrosospira* proteorhodopsin to ensure that the assembled contig did not represent a chimeric contig or inappropriate scaffolding. We verified that a single, unique path exists from the beginning to the end of the NspGL1 contig containing proteorhodopsin (Fig 3). Further, we verified that consistent coverage across this contig existed by mapping short reads from the original sample using bowtie2 (95) and viewing results using Integrated Genomics Viewer 2.7.0 (132) A closely related assembly of the same genomic region from Lake Biwa did not show evidence of proteorhodopsin; to confirm this difference between the Biwa and Great Lakes MAGs, we mapped reads from Lake Biwa (133) (BioProject PRJDB6644) onto the assembled contig described above using bowtie2 (95). This analysis demonstrated that while a large fraction of the NspGL1 contig in question recruited reads from Lake Biwa at high identity (98-99%), starting upstream of proteorhodopsin and retinal biosynthesis, this contig no longer recruited reads from Lake Biwa.

### Manual identification of key nitrification genes

Despite recovery of 15 high completion MAGs in NspGL1/2a/2b/3, many of these MAGs lacked key nitrification genes in *amo* and *hao* operons. This was largely due to the fact that *amo* and *hao* operons were often assembled on small contigs below the minimum size cutoff we imposed for binning contigs. Difficulty in assembling these contigs was likely in part due to the several *amo* and *hao* operons with extremely high identity to one another in each genome, a phenomenon which has been observed in other *Nitrosospira* (18). Manual assembly graph inspection with Bandage (131) supported this hypothesis, as did assessment of abundance of short reads associated with *amo* operons from NspGL and comparison of abundance of short reads associated with core gene *rpoB* from NspGL, using ROCker (134). Still, an exemplar MAG from at least one representative of each ecotype (NspGL1/2a/2b/3) was found with both *amo* and *hao* operons. Further, manual inspection of unbinned contigs confirmed that *amo* and *hao* operons existed on contigs in every sample from which a MAG for a particular ecotype was recovered. That is, for every time that an NspGL1 MAG was recovered from a sample, we were able to determine that an *amo* and *hao* operon which could be affiliated with NspGL1 existed, even if it was not correctly binned. Affiliation for these unbinned key nitrification genes was carried out by alignment of *amoAB* and *haoAB* sequences to *amoAB* and *haoAB* sequences correctly binned in NspGL ecotypes. This process was also carried out for two NtogaGL1a MAGs for *nxrAB*, which were poorly assembled in those two samples. Dataset S6 summarizes the presence of genes related to nitrification and nitrogen metabolism across all our MAGs.

### Verification of gene absences

Metagenome-assembled genomes typically comprise tens or even hundreds of contigs, and this fragmented nature makes it impossible to say with certainty whether a particular gene is truly absent. To substantiate our claims of gene absence based on MAGs, we used several lines of evidence. First, we note that our MAGs have high estimated completion (median 96.4%, mean 94.3%), based on the presence of universal core gene markers. Second, for all new lineages described here except NspGL3, we assembled multiple similar MAGs independently from different samples, and we inferred gene absences only if the absence was replicated in multiple assemblies. Together these two factors provide strong support for cases where a missing gene would be expected to occur in a region of predominantly core genes; however these factors are less informative for cases where a missing gene might occur in a genomic island, because we have no way of assessing the completion of regions lacking core genes, and islands tend to have systematic poor assemblies across samples. A third line of evidence that we considered is chromosome organization: if a single gene is deleted from an otherwise conserved region of synteny, then this deletion should be apparent in a gene neighborhood diagram (e.g. Figures S5, S7, S8, S10). Unfortunately in many cases, our MAGs are too dissimilar from reference genomes and share little synteny with them, so this approach is not always informative.

We used a fourth approach based on quantitative analysis of short reads to verify gene absences. If a suspected missing gene were actually present in the population, but failed to assemble and/or bin with the rest of the genome, then it should be detectable in the unassembled short reads. The frequency of a gene in the population can be estimated from its abundance in the short reads, compared to the abundance of core marker genes in the short reads. We implemented this approach as follows. We searched unassembled short reads for each gene of interest that we identified as absent from MAGs (e.g. nitrosocyanin) using tblastn. Short reads with significant similarity were then filtered by best-hit taxonomy to the appropriate nitrifier group (i.e. *Nitrosomonadaceae*, *Ca.* Nitrotoga, *Nitrospira*, *Thaumarchaea*). These filtered short reads were enumerated and length-normalized (1000 * number of short reads / length of the target gene of interest). The same procedure was repeated for genes expected to be present in every cell (e.g. *amoAB*, *hao*, *nxrAB*, ribosomal protein genes) for comparison. If a putative missing gene (based on MAGs) has near-zero detection in the short reads, we can be confident that the gene is truly missing (or has undetectable sequence similarity, or was so recently acquired from another lineage that its best hit points to a different taxon). By contrast, if a putative missing gene (based on MAGs) does recover short reads, then the gene may be present in genomes related to our MAGs but was unassembled/unbinned, or the gene may be present in another lineage of nitrifiers that is not represented by our MAGs. Short read-based quantification of select genes is presented in Supplemental Dataset S7 and described in Supplemental Text.

### Statistical analysis and plots

All statistical comparisons were carried out in R version 3.5.3 (135) and plots were generated using ggplot2 3.2.0 (136). Code and data files are available at bitbucket.org/greatlakes/gl_nitrifiers.

### Data Availability

The metagenome-assembled genomes presented here are available via NCBI BioProject PRJNA636190. 16S rRNA data is available at NCBI BioProject PRJNA591360. Metagenomes sequenced by JGI are available at http://genome.jgi.doe.gov, Project IDs 1045056, 1045059, 1045062, 1045065, 1045068, 1045071. Single-cell amplified genome is available at img.jgi.doe.gov, IMG Genome ID 3300033241. Raw reads are available in NCBI SRA (SRR14240538-SRR14240543) or through JGI with project IDs listed above.

## Acknowledgments

The work conducted by the U.S. Department of Energy Joint Genome Institute, a DOE Office of Science User Facility, is supported under Contract No. DE-AC02-05CH11231. Sequencing support was provided by the DOE JGI Community Sequencing Program (CSP #1565, #503460). Computational resources were provided by the UChicago Research Computing Center. Funding for this work was provided by Illinois-Indiana Sea Grant (Grant # NA14OAR170095), UChicago Women’s Board, and the National Science Foundation (OCE- 1830011 to MLC). We thank the science staff in the Great Lakes National Program Office of the US EPA, and the captain and crew of the R/V *Lake Guardian*, for facilitating sample collection. We thank members of the Coleman and Waldbauer labs for assistance with sample collection and processing, and for discussion and comments on the manuscript.

## Supplemental Material

**Supplemental Text.** Detailed description of short-read analyses to verify gene absences. 10.6084/m9.figshare.15130350

**Figure S1.** Relationships between environmental factors and relative abundance of nitrifiers in the Great Lakes. Relative abundance based on 16S rRNA V4-V5 region amplicon sequencing from Summer 2012. Nitrifier abundance includes sequences assigned to the families *Nitrosopumilaceae* (AOA), *Nitrosomonadaceae* (AOB), *Nitrospiraceae* (NOB), and *Gallionellaceae* (NOB). (a) Relative abundance of nitrifiers (% of total community) compared to photosynthetically active radiation (PAR). (b) Correlation between the relative abundances of nitrite oxidizers and ammonia oxidizers in each sample. A 1:1 line (solid) and linear regression fit (dashed) are shown for reference. (c) Relative abundance of nitrifiers (% of total community) compared to chlorophyll *a* concentration. PAR and chlorophyll data from the EPA Great Lakes Environmental Database (https://cdx.epa.gov/). Values are presented in Supplemental Dataset S1.

**Figure S2.** Oligotype composition of nitrifiers across the Great Lakes. Each color represents a different 16S rRNA oligotype.

**Figure S3.** Phylogenomic tree of the *Nitrosomonadales* order of *Betaproteobacteria*, focusing on the *Nitrosomonadaceae* and *Gallionellaceae* families. Tree was constructed using 139 concatenated single-copy core genes (B.J. Campbell, L. Yu, J.F. Heidelberg, and D.L. Kirchman, PNAS 108:12776-12781, 2011). Colored squares indicate the environment of origin; colored circles denote the lake of origin for genomes from the Laurentian Great Lakes; yellow stars indicate clades containing strains that lack nitrosocyanin.

**Figure S4.** Histograms of nitrifier genome properties. Rows show each taxonomic group and columns show genome properties. Each circle represents a single genome/MAG. Great Lakes MAGs shown as filled circles.

**Figure S5.** Gene neighborhood surrounding proteorhodopsin in *Nitrosospira* draft genomes from deep lakes. Light blue, shared genes on reverse strand; dark blue, shared genes on forward strand; pink, tRNA genes; orange, PR module in NspGL1 MAG; gray, genes unique to one MAG.

**Figure S6.** Phylogenomic tree of *Nitrosopumilaceae* and *Cenarchaeum*. Tree was constructed using concatenated single-copy core genes as defined in (C. Rinke *et al.,* Nature 499:431-437, 2013). Colored boxes correspond to the environment of origin; genomes from the Laurentian Great Lakes are indicated by colored circles. Names in bold indicate cultured isolates or enrichments.

**Figure S7.** Cyanase diversity and distribution among nitrite oxidizing bacteria. a) Phylogenetic tree showing distinct clades of cyanase, highlighting Great Lakes *Nitrospira* and *Ca.* Nitrotoga. b) Gene synteny surrounding the cyanase gene in NspiraGL and reference *Nitrospira*, and a cyanase insertion in NtogaGL compared to reference *Ca.* Nitrotoga.

**Figure S8.** Gene content distinguishing NtogaGL1a and NtogaGL1b. Genes colored in blue are shared between both genomes; orange colored genes are unique to either genome.

**Figure S9.** Phylogenomic tree of Lineage I and II *Nitrospira*. Tree was constructed using concatenated single-copy core genes (B.J. Campbell, L. Yu, J.F. Heidelberg, and D.L. Kirchman, PNAS 108:12776-12781, 2011). Colored boxes correspond to the environment of origin; genomes from the Laurentian Great Lakes are indicated by circles color coded by lake of origin.

**Figure S10.** Gene neighborhood surrounding photolyase genes in NspiraGL, compared to reference *N. moscoviensis*. Genes shared between the two genomes are colored light blue (protein coding) or magenta (rRNA, tRNA); non-shared genes are gray; photolyase genes are colored to match the tree in Figure 5.

**Dataset S1**. Relative abundance of nitrifiers and associated environmental parameters. Relative abundance based on 16S rRNA amplicons is reported as proportion of total community reads. The full 16S rRNA dataset and flow cytometry cell counts are presented in Paver et al. (2020) Environ Microbiol 22(1): 433-446. Chlorophyll a concentrations, irradiance values and dissolved oxygen measurements are from the US EPA Water Quality Survey (available from https://cdx.epa.gov/). Relative abundance based on metagenome read recruitment to MAGs is reported as the proportion of total sequenced reads for that sample. <u>10.6084/m9.figshare.15130350</u>

**Dataset S2**. Nitrate (NOx), ammonium, and urea concentrations compiled from literature, EPA Water Quality Survey (available from https://cdx.epa.gov/) or measured in this study. NA, not available; n.d., not detected <u>10.6084/m9.figshare.15130350</u>

**Dataset S3**. Assembly statistics and taxonomic information for metagenome-assembled genomes (MAGs) of nitrifiers in the Great Lakes. <u>10.6084/m9.figshare.15130350</u>

**Dataset S4**. Genome properties for newly assembled MAGs and reference genomes included in this study. <u>10.6084/m9.figshare.15130350</u>

**Dataset S5.** Protein functions differentiating genomes of Great Lakes nitrifiers from previously published genomes. Genome averages are calculated for reference genomes and GL MAGs. Genome-level counts for each individual GL MAG and for selected reference genomes are also shown. <u>10.6084/m9.figshare.15130350</u>

**Dataset S6.** Detailed list of presence/absence of key nitrification genes in MAGs presented in this study. NCBI Protein IDs listed when a MAG contains the target gene. 0 indicates the gene was not recovered in the MAG. <u>10.6084/m9.figshare.15130350</u>

**Dataset S7.** Short read analysis for verifying gene loss. Each target gene was searched in unassembled short reads as described in Materials & Methods. Values indicate the number of mapped reads, normalized to gene length. n.d., not detected. <u>10.6084/m9.figshare.15130350</u>

**Dataset S8.** Stations are from the US EPA Water Quality Survey (available from https://cdx.epa.gov/). <u>10.6084/m9.figshare.15130350</u>

