## Supplementary figures and images for "Genome streamlining, proteorhodopsin, and organic nitrogen metabolism in freshwater nitrifiers"

### Supplemental Figures

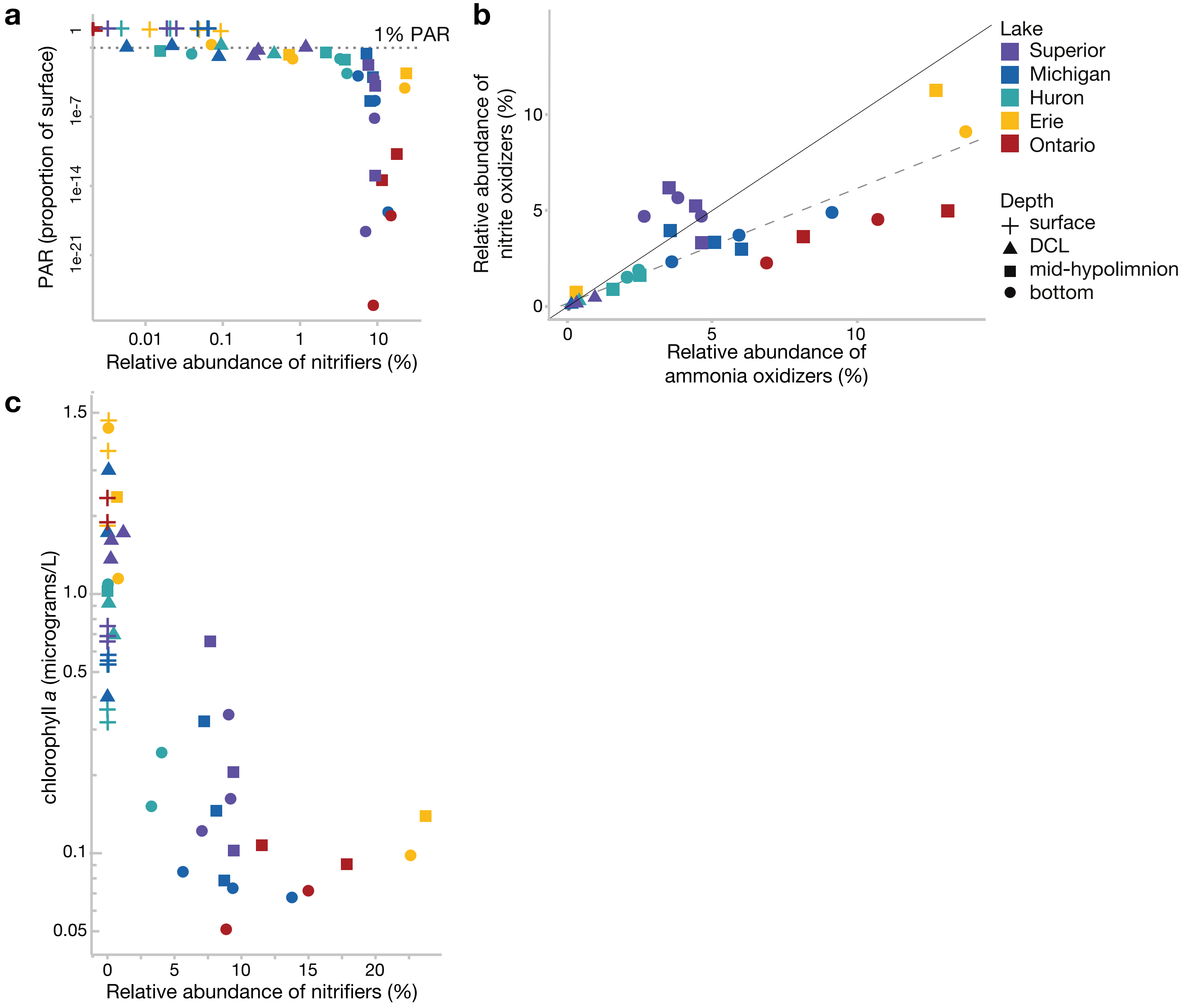

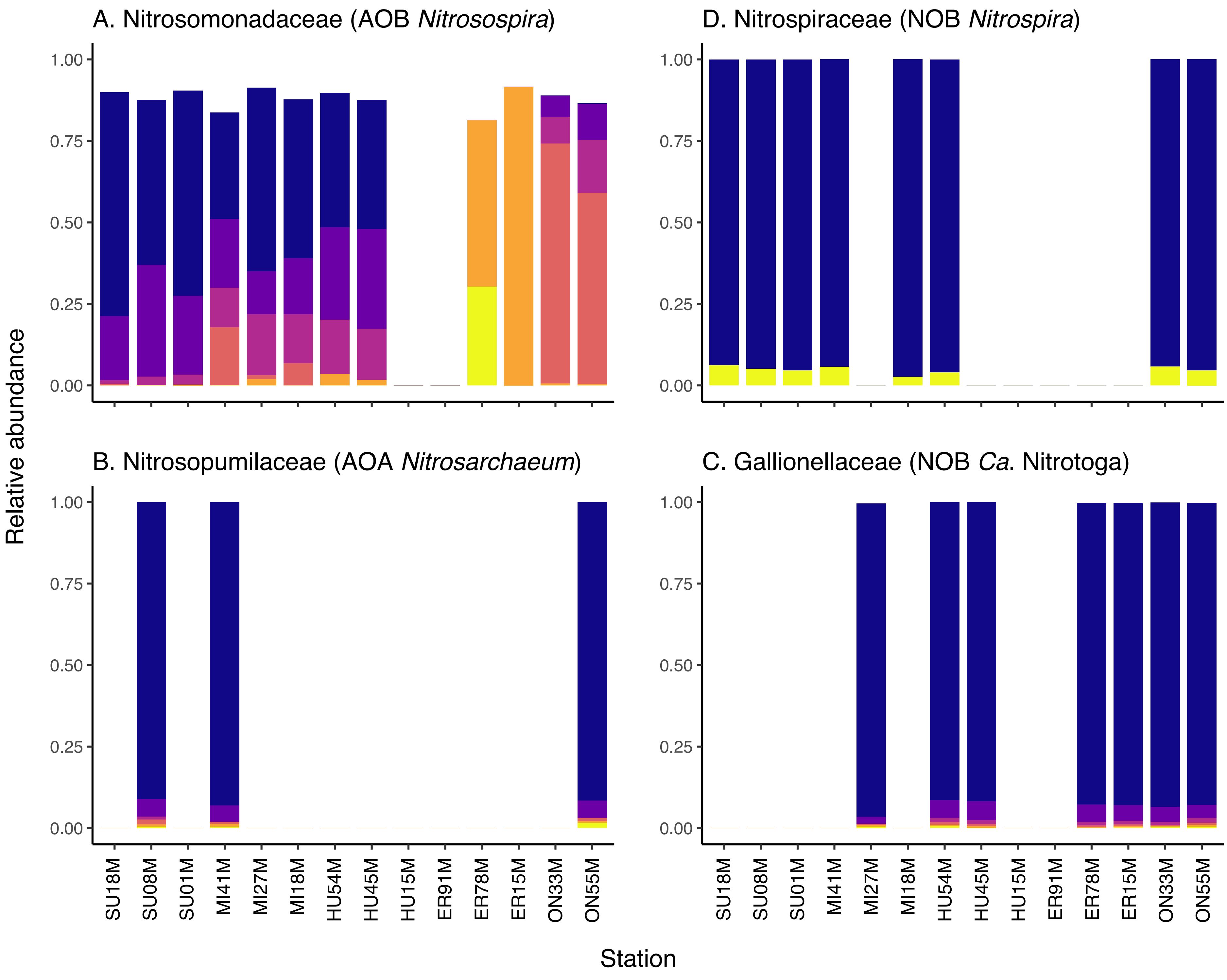

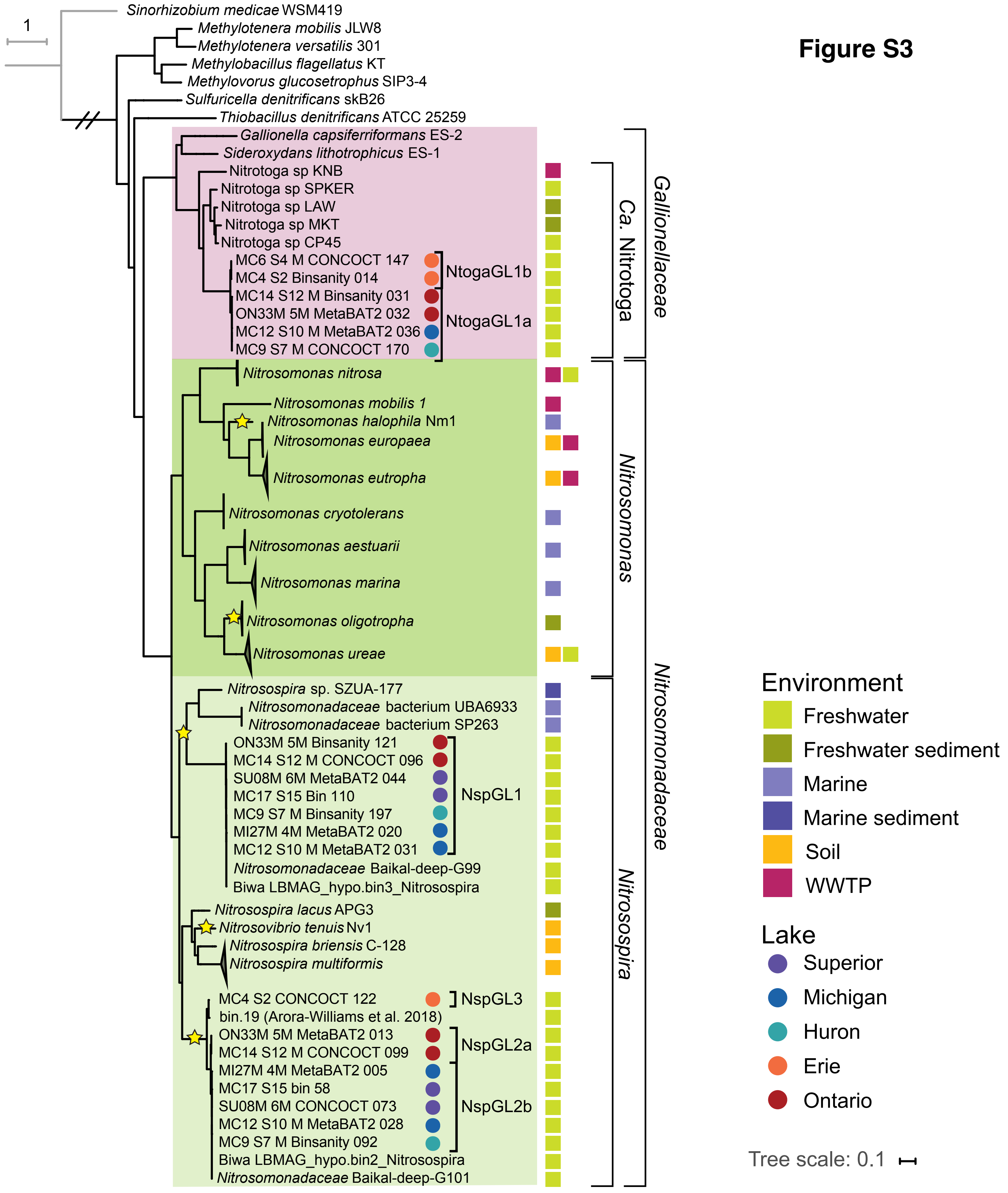

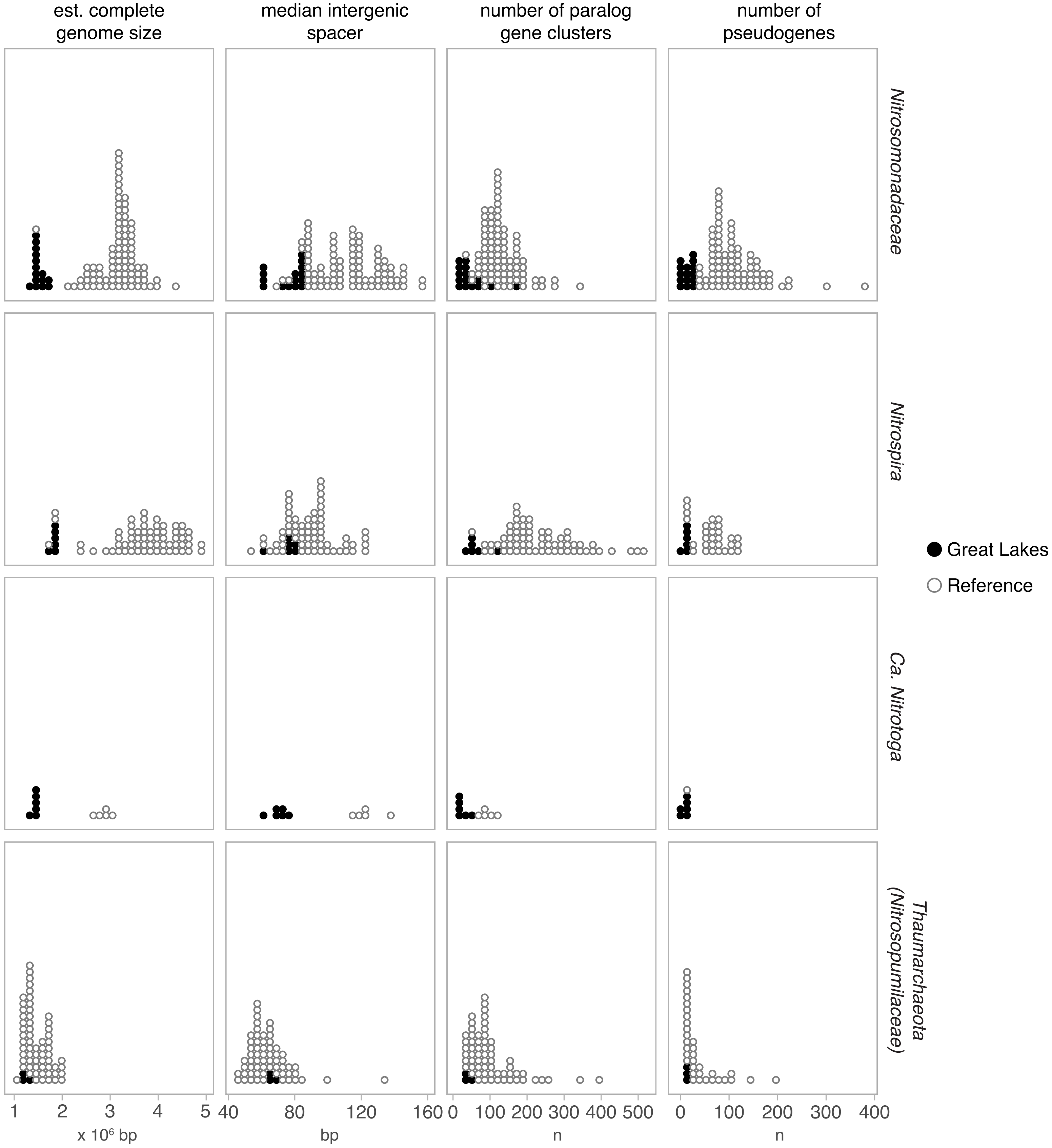

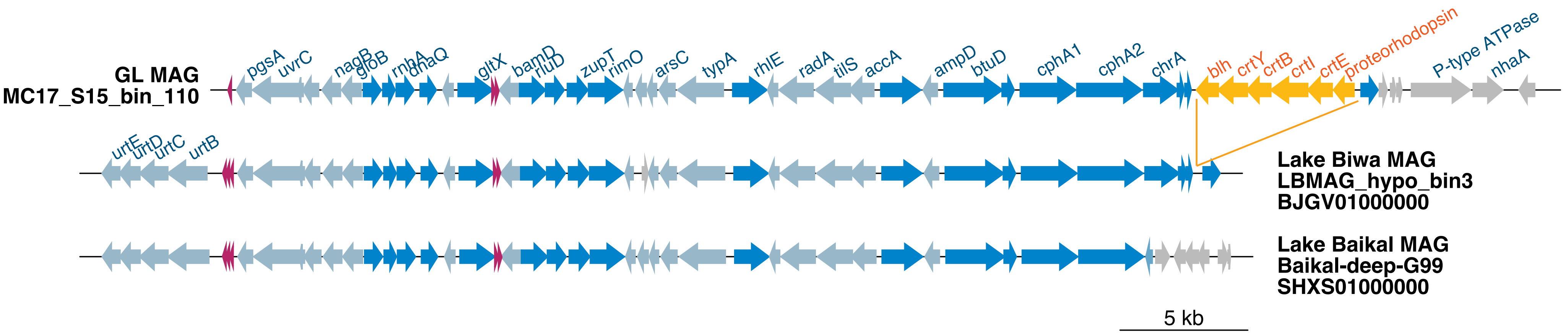

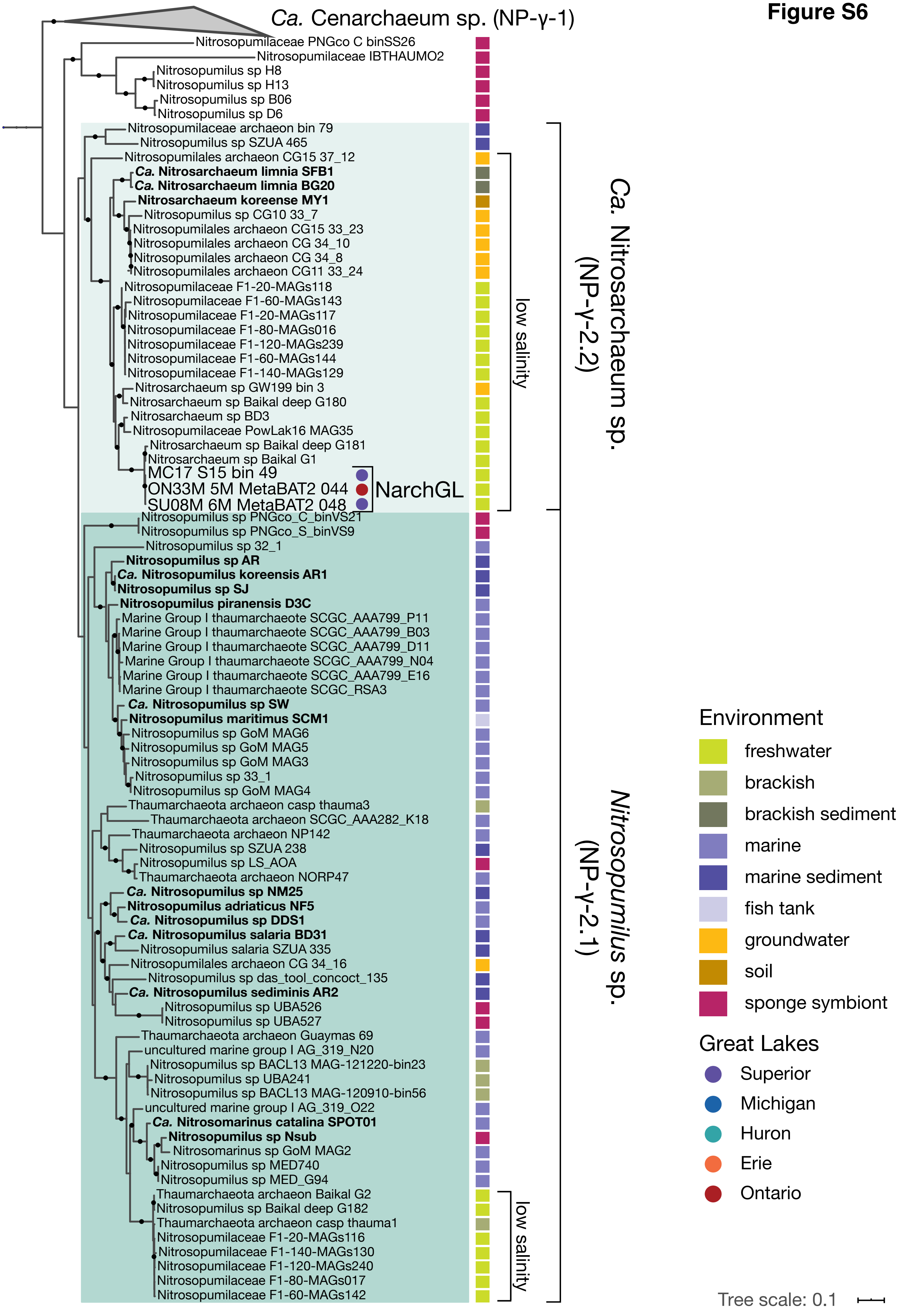

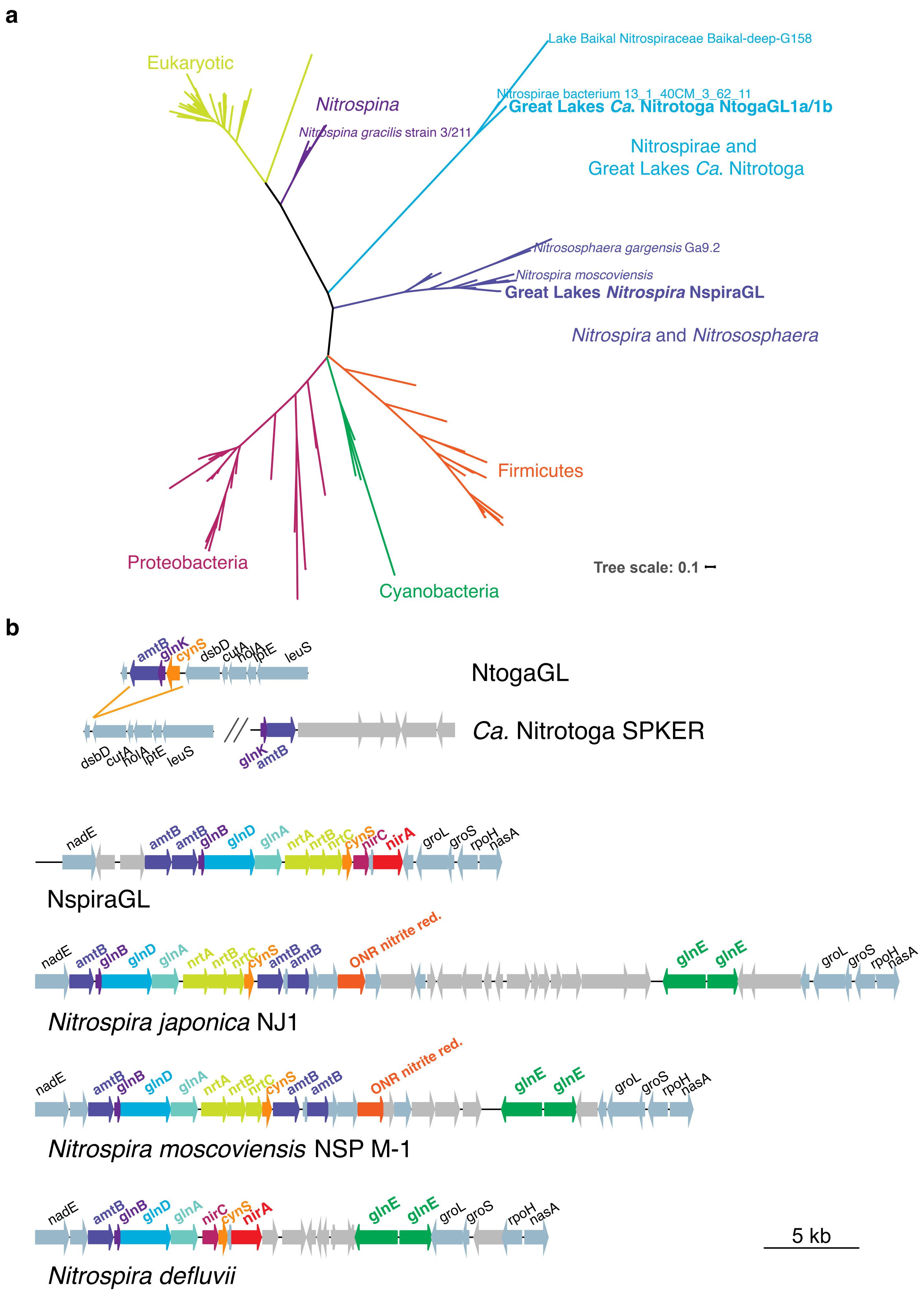

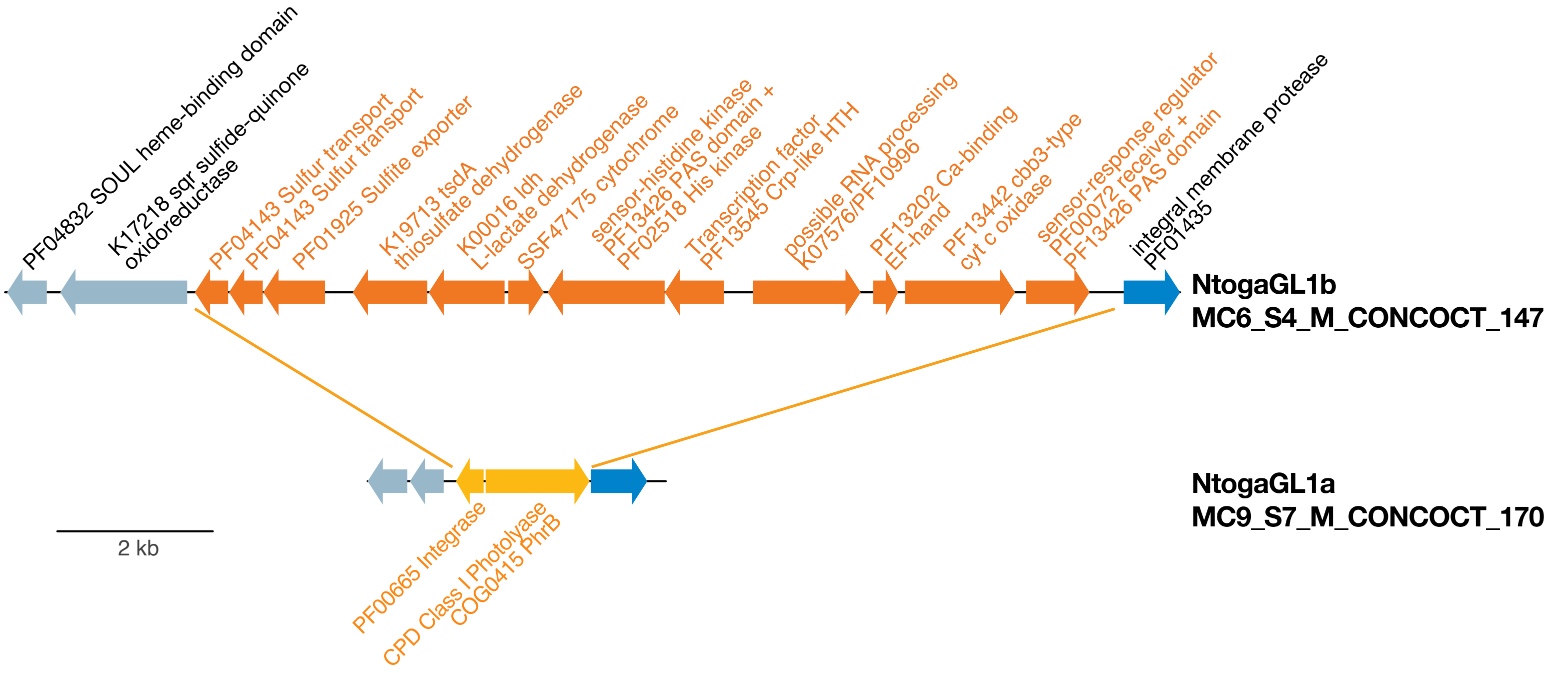

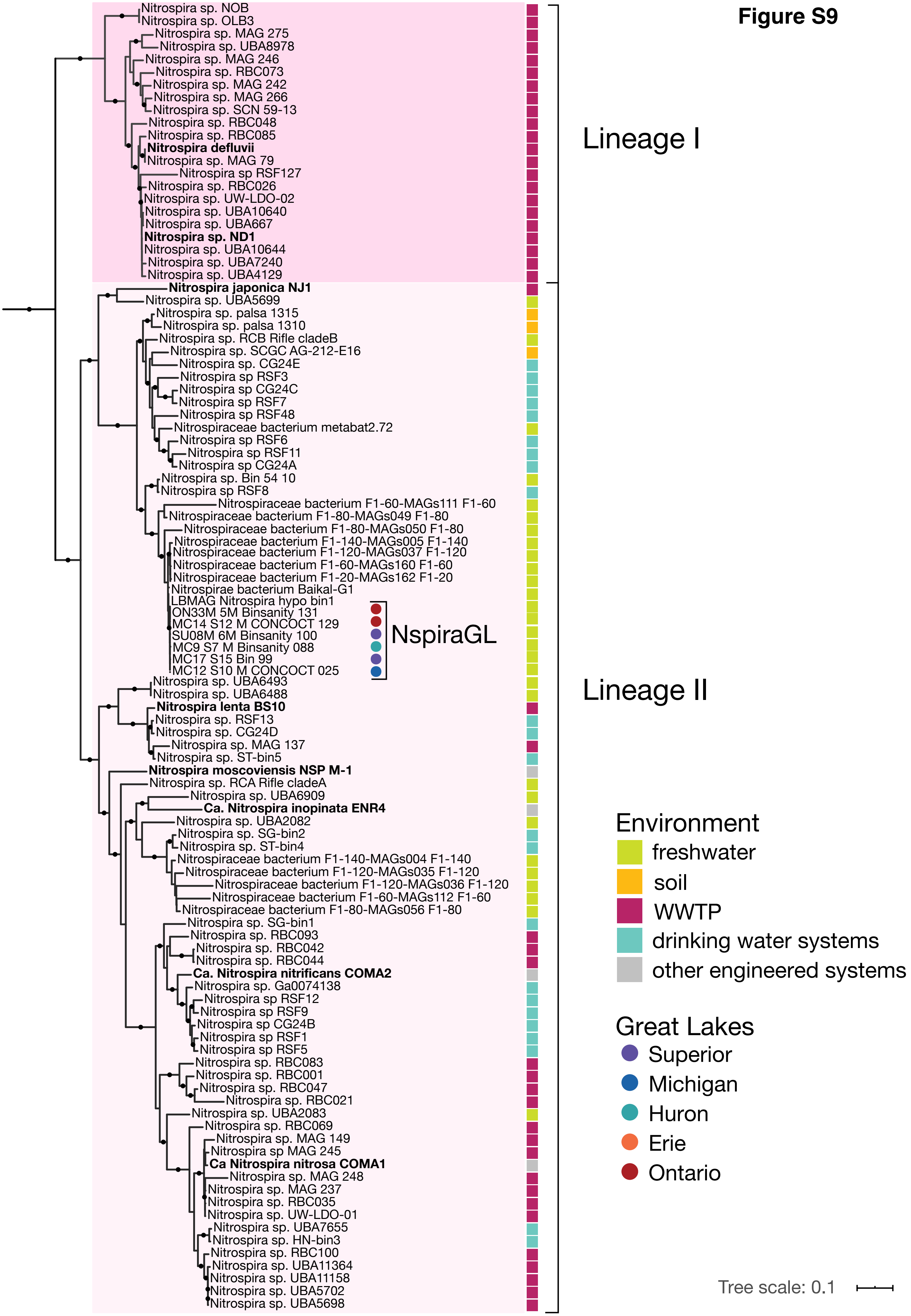

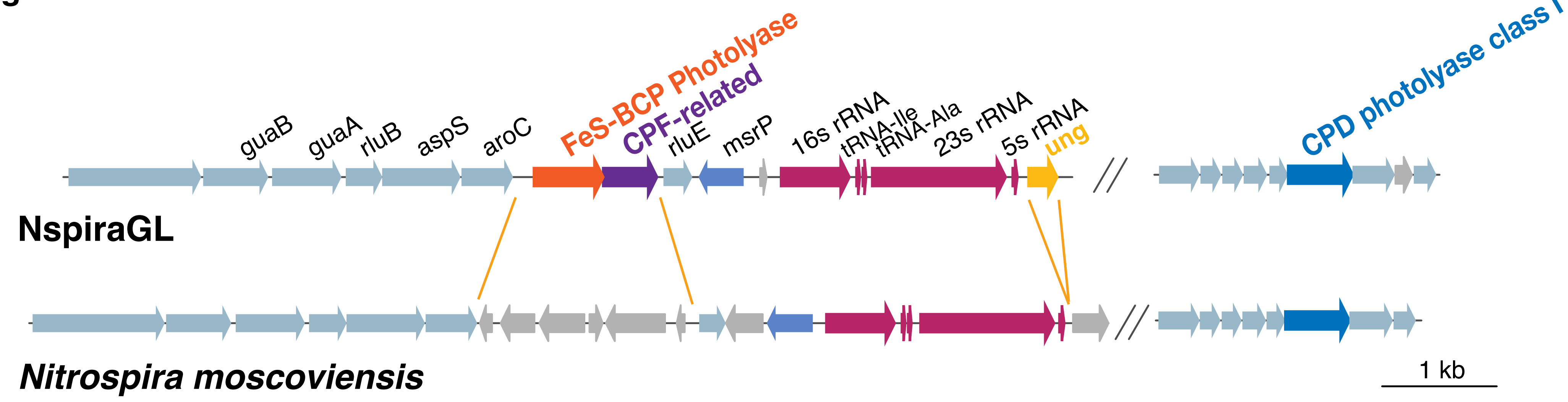
